# Habitual sleep duration is associated with the cortical organisation of brain network controllability

**DOI:** 10.64898/2026.10.25.761633

**Authors:** Xiaobo Liu, Runhao Lu, Sanwang Wang, Qiuxuan Yu, Mingyue Zhang, Ruifang Cui, Jinming Xiao, Xiaoqiang Liu, Tengteng Fan

## Abstract

Sleep is fundamental to adaptive brain function, yet how habitual sleep relates to the structural network properties that support flexible transitions between brain states remains poorly understood. Combining wrist-accelerometer sleep measures with diffusion MRI connectomes from 15,799 UK Biobank adults, we show that longer workday sleep is associated with higher modal controllability of somatomotor systems and lower controllability of visual cortex. This system-specific signature reproduced in a locked test set (n = 4,740) and was robust across extensive sensitivity analyses. Decomposition showed that it reflects a redistribution of structural connection weight, with greater absolute visual connectivity and a smaller somatomotor share of the whole-brain total. Weekend catch-up sleep showed no comparable association. The signature was distinct from brain-ageing patterns and, across 1,226 phenotypes, related mainly to physical activity and body composition rather than cognition. These findings link everyday sleep habits to the system-specific organisation of brain network controllability.

## Introduction

Sleep is fundamental to adaptive brain function, supporting the coordination of neural activity across distributed brain systems and the capacity to respond to changing cognitive and behavioural demands.^1–3^ Large population studies have linked habitual sleep duration, sleep irregularity and the balance between weekday sleep restriction and weekend recovery to cognitive performance, psychiatric disorders and mortality.^4–7^ Beyond these associations, sleep has been proposed to maintain the flexibility of brain dynamics, allowing neural systems to transition between functional states in response to internal and external demands.^1,2^ Such flexibility depends not only on local neural activity but also on how distributed brain regions interact to support transitions between different configurations of information processing. However, population-level investigations of sleep have primarily focused on regional brain morphology and functional or structural connectivity.^5,8^ Although these studies have revealed important associations between sleep and brain organisation, how habitual sleep relates to the network properties that govern the theoretical accessibility of different brain states remains poorly understood.

Network control theory provides a framework for understanding how the structural organisation of the brain supports transitions between dynamical states.^9,10^ By modelling neural activity as a dynamical system evolving on the structural connectome, this framework characterises how individual brain regions can influence the evolution of activity across the network. In particular, modal controllability quantifies the theoretical capacity of a region to drive the system towards difficult-to-reach dynamical states, providing a perspective on how structural network architecture supports a diverse repertoire of brain states.^9,10^ Previous work has shown that network controllability changes during brain development, accompanying the increasing diversity of brain dynamics, and that network control properties are relevant to the organisation of brain function in health and psychiatric disorders.^11–13^ These findings suggest that controllability may provide a useful framework for investigating the network organisation underlying adaptive brain function. Nevertheless, modal controllability remains a model-derived property rather than a direct measure of cognitive flexibility or observed state transitions. Moreover, its estimation depends strongly on the distribution and normalisation of structural connection weights, requiring careful interpretation of the biological information captured by this measure.^9,14^

These considerations raise the possibility that habitual sleep is associated with the organisation of brain network controllability across functionally distinct systems. Sleep is closely linked to neural plasticity and the regulation of distributed brain activity,^1,2^ while different cortical systems contribute to sensory processing, motor control and higher-order cognitive functions. Rather than exhibiting a uniform association with network controllability, habitual sleep may therefore be related to distinct control properties across these systems. Such system-specific organisation could provide insight into how everyday sleep habits are associated with structural network properties relevant to flexible brain-state transitions. However, several questions remain unresolved. Whether habitual sleep duration, sleep regularity and weekend recovery show distinct associations with regional and large-scale network controllability is unknown. It is also unclear whether such associations reflect changes in absolute structural connectivity or a redistribution of connection weights across functional systems, and whether they can be distinguished from broader patterns of brain ageing and genetic liability. Finally, although sleep is closely intertwined with physical activity, body composition and cognitive performance, the behavioural relevance of sleep-associated network controllability patterns remains to be established. Addressing these questions requires large-scale population data, independent replication and systematic evaluation of alternative explanations.^15^

Here, we combined seven-day wrist accelerometry with diffusion MRI connectomes from 15,799 UK Biobank adults to investigate how habitual sleep relates to the system-specific organisation of brain network controllability. We first characterised associations between multiple dimensions of habitual sleep and regional and network-level modal controllability, identifying their spatial distribution across cortical and subcortical systems. We then examined the underlying distribution of structural connection weights to determine the extent to which the observed controllability patterns reflected absolute connectivity or relative network organisation. To establish reproducibility, we evaluated a discovery-defined signature in a predefined, locked test set and assessed its robustness across imaging sites, health conditions and methodological specifications. Finally, we investigated the relationship of sleep-associated controllability patterns with brain ageing, genetic liability and 1,226 behavioural and physiological phenotypes, including cognition, physical activity and body composition (Fig. 1). Together, this approach aimed to characterise how everyday sleep habits are associated with the organisation of brain network controllability and to clarify the potential relevance of these associations to adaptive brain function.

**Fig. 1.**
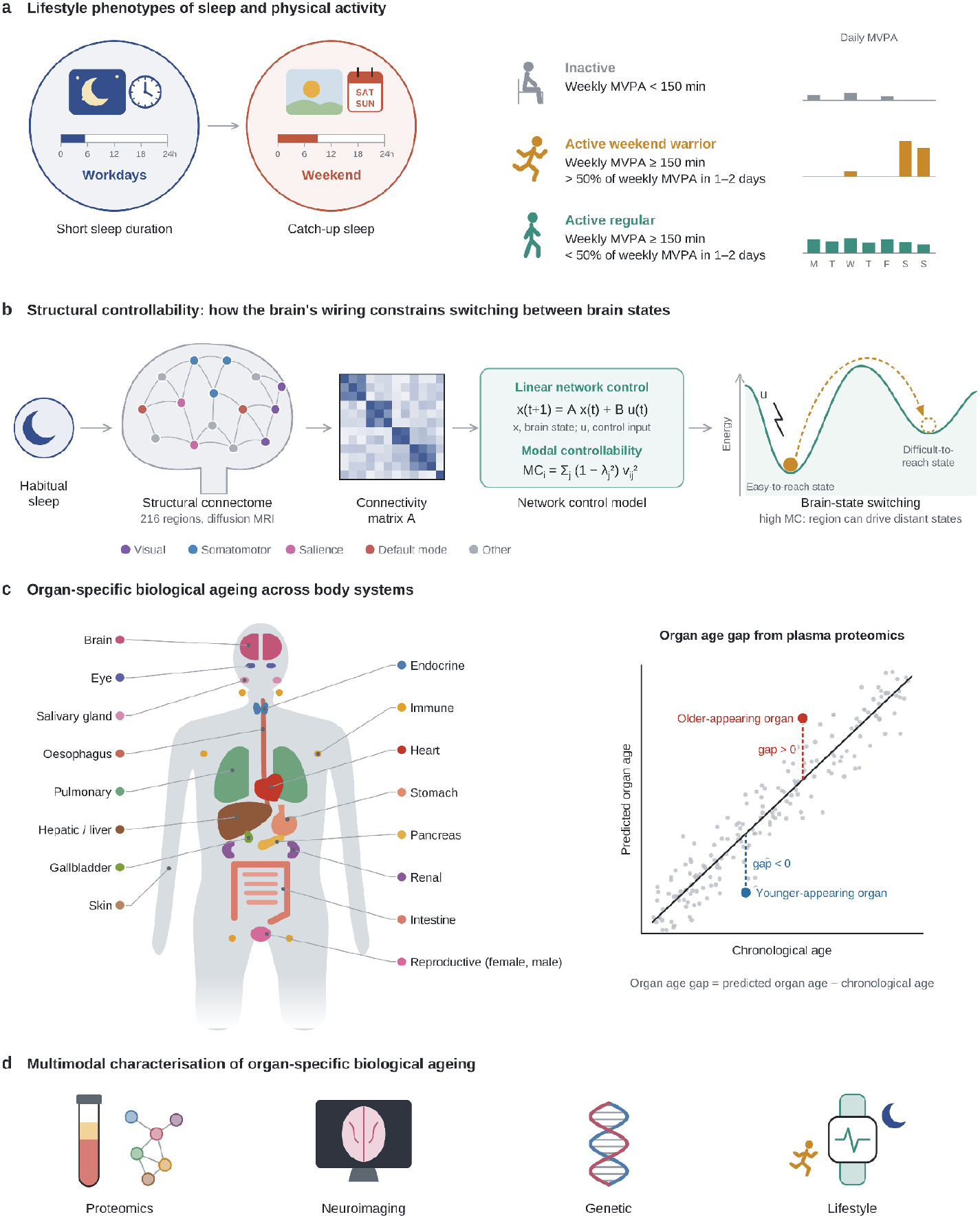
| Study design and measures. **a**, Lifestyle phenotypes from 7-day wrist accelerometry (n = 15,799): workday and weekend sleep duration, catch-up sleep (weekend minus workday sleep), and physical-activity patterns defined by weekly moderate-to-vigorous physical activity (MVPA): inactive, < 150 min per week (n = 4,734); weekend warrior, ≥ 150 min per week with > 50% accumulated in 1–2 days (n = 7,322); regular, ≥ 150 min per week otherwise (n = 3,743). Bars illustrate daily MVPA. **b**, Structural controllability. Diffusion-MRI connectomes (200 cortical and 16 deep-nuclear regions) define the connectivity matrix A of a linear network control model, x(t + 1) = Ax(t) + Bu(t). The modal controllability (MC) of region i, MC_i_ = Σ_j_(1 − λ_j_²)v_ij_², indexes its capacity to drive the system into difficult-to-reach states. **c**, Organ-specific biological ageing. Organ age was predicted from baseline plasma proteomics for 11 organ systems and 7 gastrointestinal organs (an organismal clock was also derived; Methods); the organ age gap is predicted minus chronological age (positive, older-appearing organ). **d**, Multimodal characterisation: proteomics, neuroimaging (MRI brain-age gap), polygenic scores and lifestyle measures.

## Results

### Longer habitual sleep is associated with a sensorimotor-centred controllability signature

We first asked whether habitual sleep is associated with regional modal controllability (MC) at all and, if so, which systems carry the association. We studied 15,799 UK Biobank participants (mean age 64.1 years; 55.2% women) with 7-day wrist accelerometry and a diffusion connectome at the first imaging visit (Table 1 and Supplementary Fig. 1).^16–19^ The connectome comprised 200 cortical and 16 deep-nuclear nodes.^20,21^ Before any revision analysis, participants were split once, stratified by site, into a discovery set (70%, n = 11,059) and a locked test set (30%, n = 4,740). Unless stated otherwise, estimates are from ordinary least-squares regression of z-scored MC on each sleep measure with the full covariate set (Methods); β is the difference in MC (s.d.) per additional hour of sleep, with 95% confidence intervals (CI), two-sided P values and Benjamini–Hochberg false discovery rate (q).

**Table 1.** | Characteristics of the analytic sample.

| **Characteristic** | **Total** | **Discovery (70%)** | **Locked test (30%)** |
| --- | --- | --- | --- |
| Participants, n | 15,799 | 11,059 | 4,740 |
| Age at imaging, years | 64.1 (7.7) | 64.1 (7.7) | 64.1 (7.8) |
| Women, n (%) | 8,723 (55.2) | 6,124 (55.4) | 2,599 (54.8) |
| Body mass index, kg m<sup>-2</sup> | 26.2 (4.3) | 26.2 (4.3) | 26.2 (4.2) |
| Townsend deprivation index | -1.92 (2.71) | -1.93 (2.71) | -1.90 (2.73) |
| College or university degree, n (%) | 7,623 (48.7) | 5,348 (48.8) | 2,275 (48.3) |
| White ethnicity, n (%) | 15,372 (97.5) | 10,755 (97.5) | 4,617 (97.5) |
| Workday sleep, h | 8.64 (1.19) | 8.64 (1.20) | 8.63 (1.19) |
| Weekend sleep, h | 9.19 (1.42) | 9.17 (1.41) | 9.21 (1.42) |
| Catch-up sleep, h | 0.55 (1.26) | 0.53 (1.26) | 0.58 (1.26) |
| Sleep irregularity, h | 1.17 (0.57) | 1.17 (0.56) | 1.18 (0.58) |
| MVPA, min per week, median (IQR) | 259 (130–432) | 259 (130–432) | 259 (130–418) |
| Physical-activity pattern |  |  |  |
| Inactive, n (%) | 4,734 (30.0) | 3,283 (29.7) | 1,451 (30.6) |
| Weekend warrior, n (%) | 7,322 (46.3) | 5,189 (46.9) | 2,133 (45.0) |
| Regular, n (%) | 3,743 (23.7) | 2,587 (23.4) | 1,156 (24.4) |
| Accelerometry-to-scan interval, years | 2.99 (1.77) | 2.97 (1.78) | 3.03 (1.75) |
| Imaging site |  |  |  |
| Cheadle, n (%) | 9,237 (58.5) | 6,466 (58.5) | 2,771 (58.5) |
| Reading, n (%) | 2,625 (16.6) | 1,837 (16.6) | 788 (16.6) |
| Newcastle, n (%) | 3,917 (24.8) | 2,742 (24.8) | 1,175 (24.8) |
| Bristol, n (%) | 20 (0.1) | 14 (0.1) | 6 (0.1) |

Values are mean (s.d.) unless stated otherwise. The site-stratified split was fixed before revision analyses. MVPA, moderate-to-vigorous physical activity. Activity patterns: inactive, <150 min per week of MVPA; weekend warrior, ≥150 min per week with ≥50% in 1–2 days; regular, otherwise.

In the discovery set, longer workday sleep was associated with regional MC in 15 of 216 nodes after FDR correction across nodes (|t| = 2.93–3.94, all P ≤ 0.003; 11 positive and 4 negative; Fig. 2a). Longer sleep was associated with higher MC in medial prefrontal and medial parietal default-mode, orbitofrontal, ventral prefrontal and somatomotor cortex. It was associated with lower MC in visual cortex and left lateral prefrontal control regions. At the network level, three associations survived FDR correction pooled across six sleep measures and ten network outcomes (Fig. 2b, Table 2 and Supplementary Fig. 3).^22^ These were longer workday sleep with higher somatomotor MC (β = 0.025 s.d. per hour, 95% CI 0.012 to 0.038, P = 1.2 × 10⁻⁴, q = 0.007, n = 15,799); longer workday sleep with lower visual MC (β = −0.018, 95% CI −0.030 to −0.006, P = 0.002, q = 0.049); and longer weekend sleep with higher somatomotor MC (β = 0.019, 95% CI 0.008 to 0.029, P = 6.7 × 10⁻⁴, q = 0.020). Weekend sleep was also nominally associated with lower visual MC (β = −0.012, 95% CI −0.022 to −0.003, P = 0.012, q = 0.17).

**Fig. 2.**
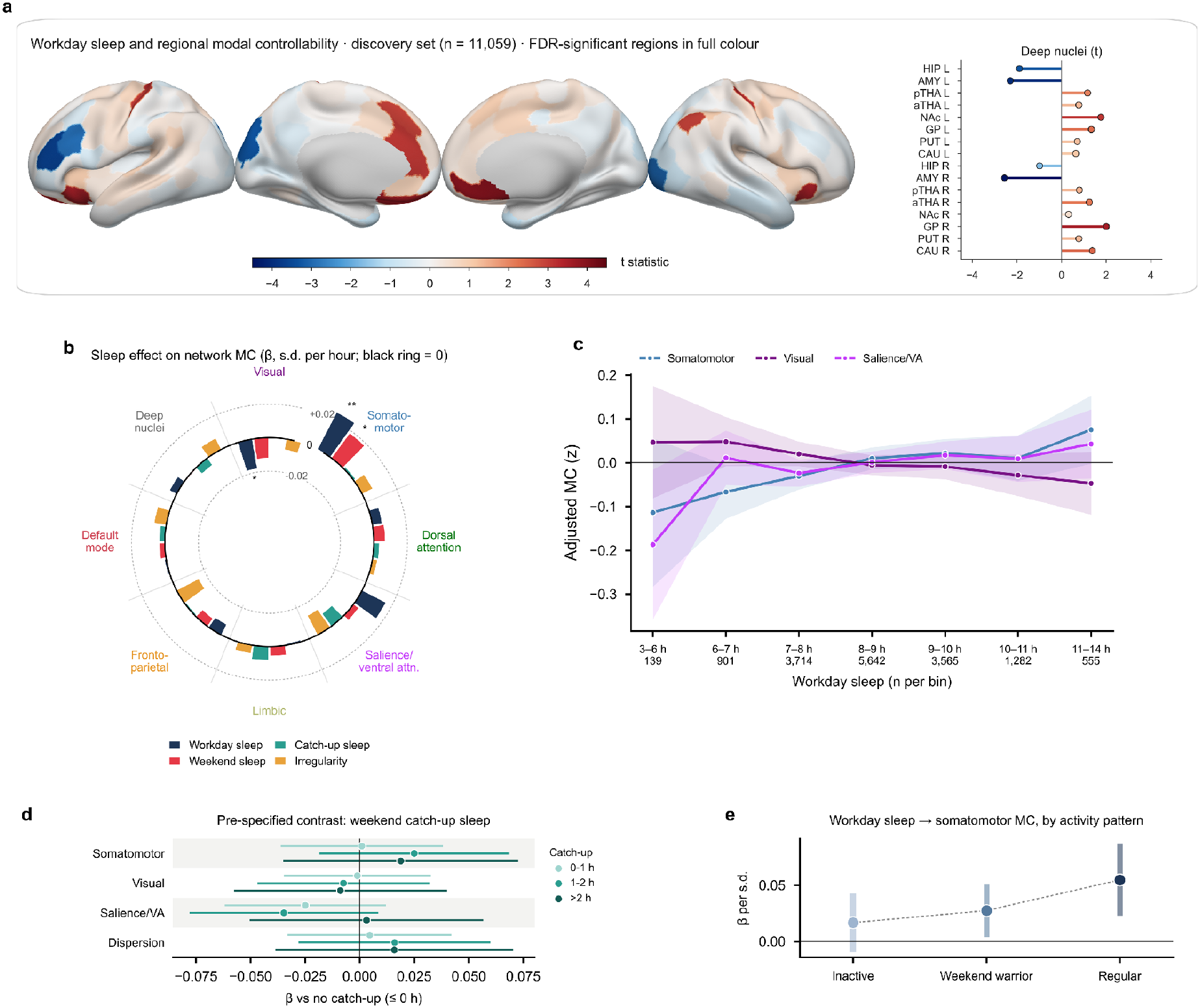
| Longer habitual sleep is associated with a sensorimotor-centred controllability signature. **a**, Discovery-set (n = 11,059) regional t-statistics for workday sleep and modal controllability (MC); non-significant regions are shown at 35% intensity; lollipops show the 16 deep nuclei. **b**, Circular barplot of the sleep effect on network MC in the full sample (n = 15,799). Each sector is a network (deep nuclei pooled); each bar is β, the change in MC (in s.d.) per additional hour of sleep (per s.d. for irregularity), drawn outward (positive) or inward (negative) from the black zero ring; dotted rings mark ±0.02. Stars denote FDR pooled across exposures (one star, q < 0.05; two stars, q < 0.01). Colour bar in **a**: regional t-statistic. **c**, Covariate-adjusted network MC across workday-sleep bins (line, mean; ribbon, 95% CI; n per bin under each label). **d**, Weekend catch-up sleep groups versus no catch-up. **e**, Workday sleep–somatomotor MC association within physical-activity patterns.

**Table 2.** | Habitual sleep and network modal controllability in the full sample, the discovery set and the locked test set.

| Exposure | Network MC | Full sample $\beta$ [95% CI] | q | Discovery $\beta$ [95% CI] | Locked test $\beta$ [95% CI] | P (test) |
| --- | --- | --- | --- | --- | --- | --- |
| Workday sleep | Somatomotor | 0.025 [0.012, 0.038] | 0.007 | 0.026 [0.011, 0.041] | 0.024 [0.000, 0.047] | 0.05 |
|  | Visual | -0.018 [-0.030, -0.006] | 0.05 | -0.022 [-0.036, -0.008] | -0.008 [-0.030, 0.013] | 0.44 |
|  | Salience/ventral attention | 0.016 [0.003, 0.029] | 0.17 | 0.021 [0.006, 0.036] | 0.005 [-0.019, 0.028] | 0.69 |
|  | Dispersion | 0.014 [0.001, 0.027] | 0.28 | 0.014 [-0.002, 0.029] | 0.015 [-0.008, 0.039] | 0.20 |
| Weekend sleep | Somatomotor | 0.019 [0.008, 0.029] | 0.02 | 0.022 [0.010, 0.035] | 0.010 [-0.010, 0.029] | 0.34 |
|  | Visual | -0.012 [-0.022, -0.003] | 0.17 | -0.019 [-0.030, -0.007] | 0.003 [-0.015, 0.020] | 0.78 |
|  | Salience/ventral attention | 0.004 [-0.007, 0.015] | 0.87 | 0.011 [-0.002, 0.023] | -0.014 [-0.034, 0.005] | 0.16 |
|  | Dispersion | 0.011 [0.000, 0.022] | 0.30 | 0.009 [-0.004, 0.022] | 0.017 [-0.003, 0.037] | 0.09 |

$\beta$ is the difference in modal controllability (MC; s.d.) per additional hour of sleep from linear models adjusted for age, age<sup>2</sup>, sex, age × sex, site, intracranial volume, diffusion motion, scanner position, accelerometry-to-scan interval, Townsend index, body mass index, education, smoking, alcohol, ethnicity and physical activity. q, false discovery rate across six sleep measures and ten network outcomes (full sample). Full sample n = 15,799; discovery n = 11,059; test n = 4,740.

The relationship was graded across the sleep distribution, with no evidence of non-linearity (likelihood-ratio test of a cubic regression spline (4 d.f.) against a linear term: somatomotor P = 0.50, visual P = 0.95, salience/ventral attention P = 0.43; Fig. 2c). Weekend catch-up sleep was pre-specified as a contrast that indexes compensation rather than habitual duration. It was not associated with MC, whether analysed continuously (somatomotor β = 0.001, 95% CI −0.011 to 0.014, P = 0.82; visual β = 0.000, −0.011 to 0.011, P = 0.98) or categorically against no catch-up (all P ≥ 0.26; trend P = 0.28 and 0.65; Fig. 2d). The somatomotor association was positive in all three activity patterns: inactive participants (β = 0.017 per s.d., 95% CI −0.009 to 0.043, n = 4,734), weekend warriors (β = 0.027, 0.004 to 0.051, n = 7,322) and regular exercisers (β = 0.055, 0.023 to 0.087, n = 3,743; Fig. 2e and Supplementary Fig. 6).^23^

### The signature reflects a redistribution of connection weight across sensory systems

We next asked what the signature reflects in terms of connection weight: a change in the absolute connectivity of the implicated systems or a redistribution of connection weight across systems, and whether it is independent of how strongly connected a region is. Group-average MC was highest in somatomotor and salience cortex and lowest in the strongly connected deep nuclei (Fig. 3a). This is consistent with the inverse relation between modal controllability and connection strength. In the linear model,^9,10^ MC_i_ = Σ_j_(1 − λ_j_²)v_ij_², which for orthonormal eigenvectors reduces to MC_i_ = 1 − d_i_/c². Here d_i_ = Σ◻ A_i_◻² is the node’s squared strength and c is the stabilising constant. With the available normalisation, MC is therefore an exact monotone function of each node’s share of whole-brain squared connectivity; the rank correlation within each participant was −1.000 (Fig. 3b). We therefore decomposed the sleep association (Fig. 3c). Longer sleep was associated with greater absolute squared connectivity of visual cortex (β = 0.016, 95% CI 0.006 to 0.027, P = 0.002, q = 0.017). Somatomotor connectivity showed no absolute difference (β = −0.004, 95% CI −0.015 to 0.007, P = 0.49) but formed a smaller share of the whole-brain total (β = −0.025, 95% CI −0.038 to −0.012, P = 1.2 × 10⁻⁴, q = 0.003); whole-brain squared connectivity itself did not differ (β = 0.010, 95% CI −0.001 to 0.020, P = 0.064). Sleep-related differences were modestly larger in more strongly connected nodes (regional Pearson correlation with hubness r = −0.28, parametric P = 3 × 10⁻⁵, 216 nodes; Fig. 3d). A multivariate partial least squares (PLS) analysis^24^ of six sleep features against 216 nodes yielded one significant latent variable (latent correlation r = 0.105, 49.0% of cross-block covariance, permutation P = 0.002; LV2 and LV3, P = 0.21 and 0.10). This latent variable was dominated by workday and weekend sleep duration and involved pallidal and putaminal nuclei (Fig. 3e).

**Fig. 3.**
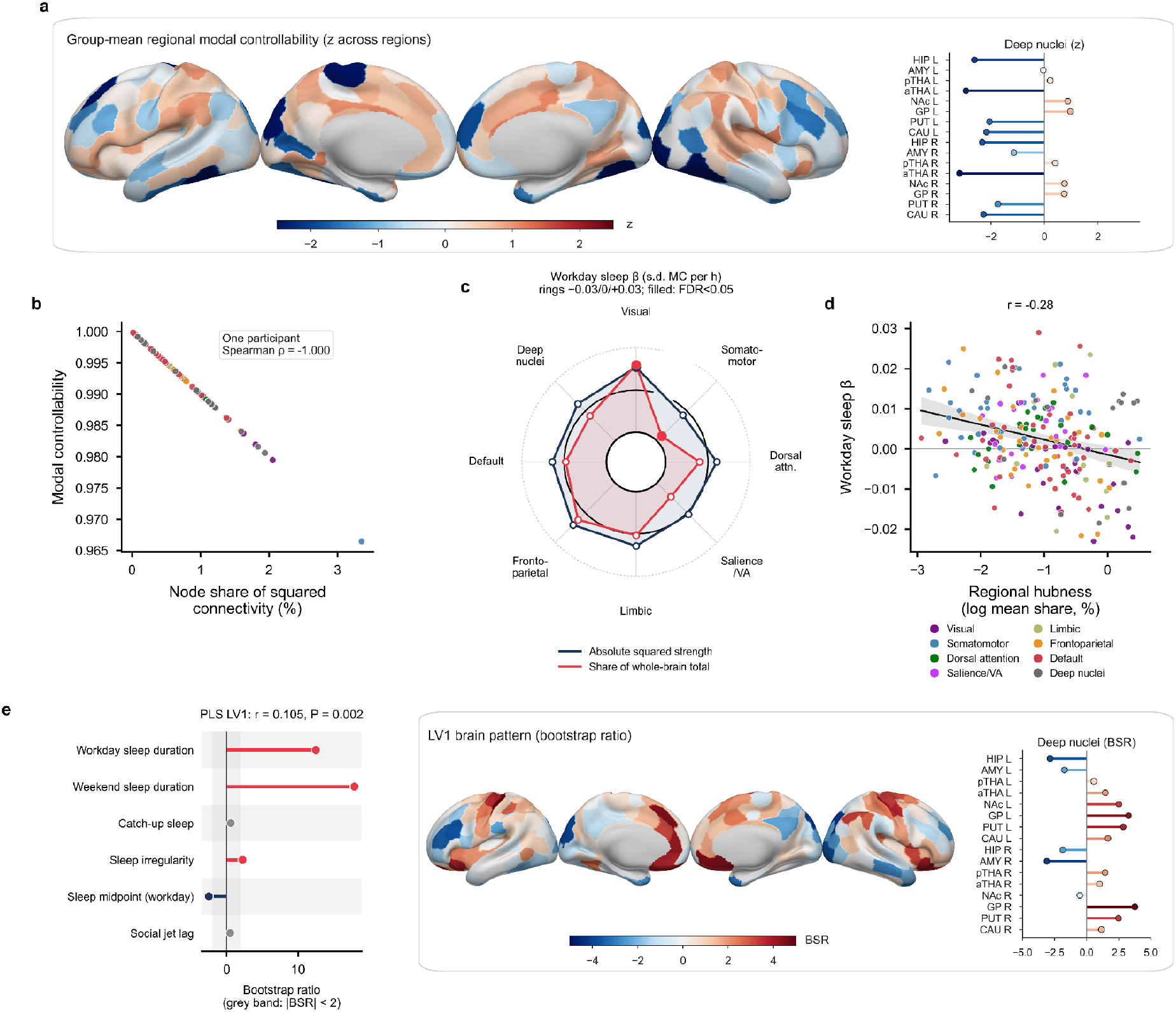
| The signature reflects a redistribution of connection weight. **a**, Group-mean regional MC (z across regions). **b**, Identity between MC and node share of squared connectivity in one participant. **c**, Radar decomposition of the workday-sleep association (β, s.d. per hour) into absolute network squared strength (blue) and share of the whole-brain total (red); the bold ring is β = 0 and the dotted rings ±0.03; filled markers, FDR < 0.05. **d**, Regional sleep effect versus regional hubness. **e**, PLS latent variable 1: sleep bootstrap ratios (left) and node bootstrap ratios (right).

### The signature is reproducible and specific

To establish reproducibility, we asked whether the discovery-defined signature reproduces in participants never used to define it, and whether disease, health, imaging quality, identifier linkage, imaging site or analytic choices explain it. Regional effect maps estimated independently in the discovery and test sets were correlated (Pearson r = 0.39 across 216 nodes, permutation P = 0.001, 1,000 permutations of sleep within the test set), and 14 of 15 (93%) discovery-significant nodes kept their sign (Fig. 4a). A signature score defined in the discovery set was associated with workday sleep in the untouched test set (β = 0.047 s.d. per hour, 95% CI 0.022 to 0.071, P = 2.0 × 10⁻⁴, n = 4,740), with weekend sleep (β = 0.023, 95% CI 0.003 to 0.044, P = 0.024) and with irregularity (β = 0.073, 95% CI 0.023 to 0.123, P = 0.005). It was not associated with catch-up sleep (β = −0.011, 95% CI −0.034 to 0.012, P = 0.35) or social jet lag (β = 0.019, 95% CI −0.011 to 0.049, P = 0.22; Fig. 4b). Connectome identifiers had to be recovered by sequence alignment (Methods). Estimates were unchanged in participants mapped without ambiguity (n = 8,799; somatomotor β = 0.025, 95% CI 0.007 to 0.042, P = 0.005; visual β = −0.020, 95% CI −0.035 to −0.004, P = 0.013) and in subsets with alignment margins > 2 (n = 10,989) or > 4 (n = 9,673; somatomotor β = 0.022–0.025, visual β = −0.020; all P ≤ 0.008; Fig. 4c and Supplementary Fig. 2).

**Fig. 4.**
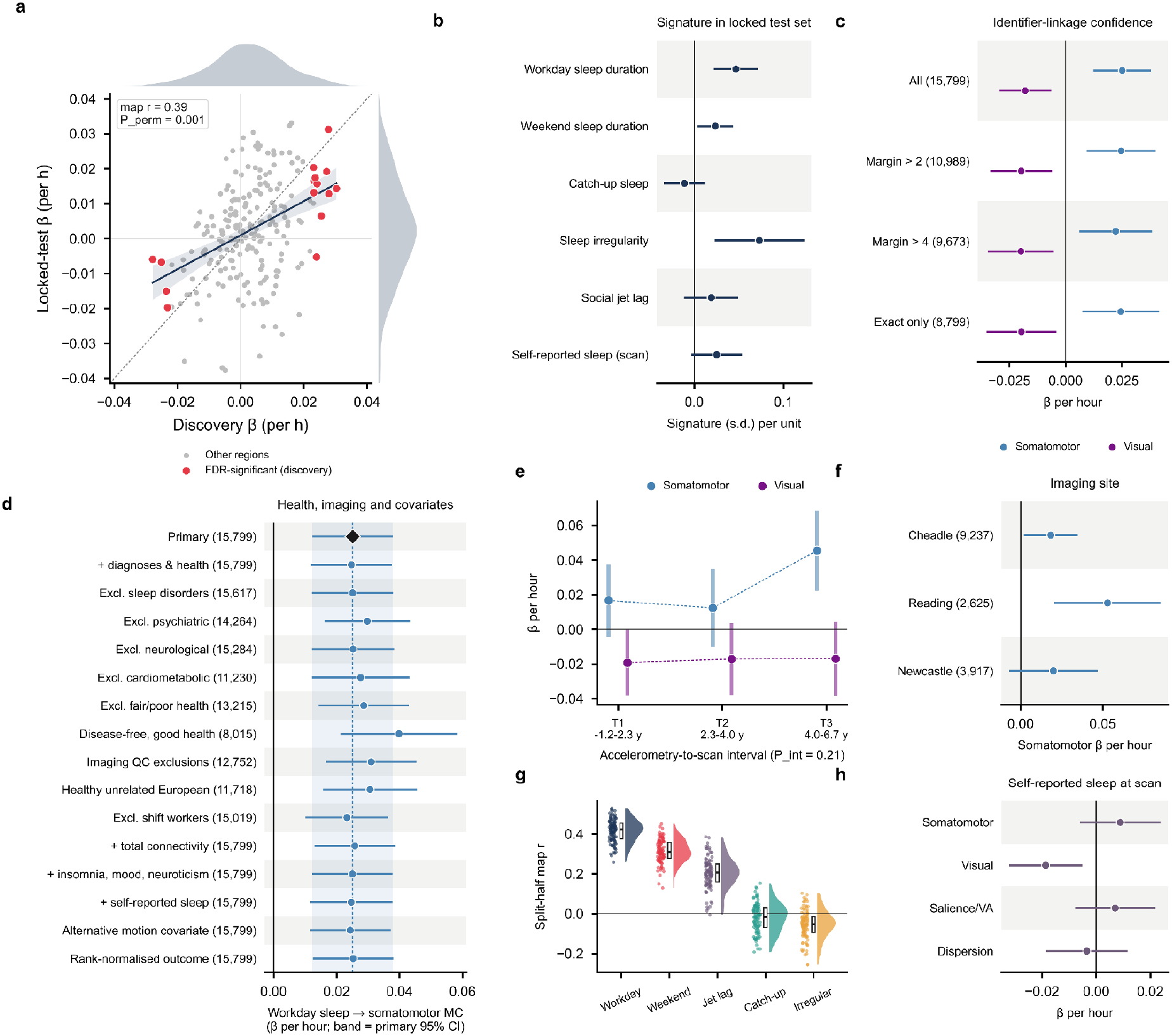
| The signature is reproducible and specific. **a**, Discovery versus locked-test regional β (permutation test of map correlation). **b**, Discovery-defined signature in the test set (n = 4,740). **c**, High-confidence identifier subsets. **d**, Disease, health, imaging-quality and covariate analyses (n in brackets; diamond, dashed line and shaded band, primary estimate and its 95% CI). **e**, Accelerometry-to-scan interval tertiles. **f**, Imaging sites (Bristol, n = 20, not estimable). **g**, Split-half map correlations (100 splits; raincloud plots show the density, individual splits and the median with interquartile range). **h**, Self-reported sleep at the imaging visit.

The somatomotor association survived adjustment for, or exclusion of, sleep, psychiatric, neurological and cardiometabolic disease. It also survived adjustment for self-rated health and long-standing illness, exclusion of low-quality imaging, and adjustment for total connectivity, self-reported sleep, insomnia, mood and neuroticism (β = 0.023–0.040; all P ≤ 5 × 10⁻⁴; Fig. 4d and Supplementary Fig. 4). It did not differ by the accelerometry-to-scan interval (interval × sleep interaction P = 0.21; tertile estimates β = 0.017, 0.012 and 0.045; Fig. 4e). It was positive at each imaging site with an estimable sample (Cheadle β = 0.018, 95% CI 0.002 to 0.035, n = 9,237; Reading β = 0.053, 95% CI 0.021 to 0.085, n = 2,625; Newcastle β = 0.020, 95% CI −0.007 to 0.047, n = 3,917; Fig. 4f and Supplementary Fig. 5). Regional maps were reproducible across 100 random split halves for workday sleep (median r = 0.42, 2.5th–97.5th percentile 0.30 to 0.52) and weekend sleep (median r = 0.31, 0.20 to 0.42), but not for catch-up sleep (median r = −0.02, −0.17 to 0.12) or irregularity (median r = −0.05, −0.20 to 0.08; Fig. 4g and Supplementary Fig. 5). Sleep self-reported at the scan replicated the visual association (β = −0.019, 95% CI −0.032 to −0.005, P = 0.007, n = 15,674) but not the somatomotor association (β = 0.009, 95% CI −0.006 to 0.024, P = 0.24; Fig. 4h).

We added three further layers of checks (Supplementary Figs. 9–11). Across 150 analytic specifications^25^ that combined five samples, five covariate sets and three outcome and two exposure treatments, every workday-sleep estimate for somatomotor MC was positive and nominally significant (median β = 0.026, range 0.017–0.038; permutation P = 0.002). Every visual estimate was negative (median β = −0.016) and 92 of 150 were nominally significant (permutation P = 0.002; Supplementary Fig. 9a,b). Robust and median regression and removal of influential observations left the somatomotor estimate unchanged (β = 0.025–0.032); the visual estimate was attenuated only under median regression (β = −0.011, P = 0.11; Supplementary Fig. 9c). Further adjustment for scan date and season, accelerometry season, snoring, napping and daytime dozing, chronotype, overall acceleration or genetic principal components changed neither estimate materially (somatomotor β = 0.022–0.029; visual β = −0.016 to −0.022; Supplementary Fig. 10a). No interaction with age, sex, BMI, chronotype, activity pattern or site survived correction (smallest P_int = 0.02 for chronotype on visual MC, 1 of 24 tests; Supplementary Fig. 10c,d). Because the standardised effects are small, the E-values^26^ were modest (1.20 for somatomotor and 1.16 for visual MC), so an unmeasured confounder associated with both sleep and MC by a risk ratio of about 1.2 could explain them (Supplementary Fig. 10b). In 1,000 bootstrap resamples the regional map was stable (median correlation with the full-sample map r = 0.85), 65 nodes kept their sign in at least 95% of resamples, and the somatomotor and visual effects kept their sign in 100% and 99.9% (Supplementary Fig. 11a,b). In subsamples, the somatomotor association reached P < 0.05 in at least 80% of draws from about 7,900 participants and the visual association from about 11,000 (Supplementary Fig. 11c). The somatomotor association held when each imaging site was left out (β = 0.020–0.033, all P ≤ 0.005), whereas the visual association was not significant without the largest site (β = −0.010, P = 0.31; Supplementary Fig. 11d). The discovery-defined signature replicated in the test set when any single network was removed (β = 0.038–0.051 per hour, all P ≤ 0.003; Supplementary Fig. 11e), so it does not depend on one system.

### The signature is distinct from brain ageing and aligned with genetic liability to affective disorders

If the signature were a marker of an older-appearing brain, its regional pattern should resemble that of brain ageing; if it shared liability with sleep-related disorders, polygenic scores for those disorders should align with it. We tested both. Across 38,742 participants with a connectome, an older-appearing brain (higher MRI brain-age gap^27^) was associated with lower cortical MC (β = −0.073 per s.d., 95% CI −0.085 to −0.061, P = 4 × 10⁻³¹, q = 1 × 10⁻²⁸) and higher deep-nuclear MC (β = 0.073, 95% CI 0.061 to 0.085, P = 4 × 10⁻³¹, q = 1 × 10⁻²⁸; Fig. 5a). The sleep map was not correlated with this ageing map (Pearson r = −0.01 across 216 nodes, parametric P = 0.88; Fig. 5b). Within the same 4,979 participants, a proteomic brain-age gap^28,29^ measured nine years earlier and the MRI brain-age gap were nearly uncorrelated (r = 0.09, P = 3 × 10⁻¹⁰). When entered together, they showed opposite associations with salience/ventral-attention MC (proteomic β = 0.064, 95% CI 0.037 to 0.091, P = 4 × 10⁻⁶; MRI β = −0.060, 95% CI −0.100 to −0.019, P = 0.004; Fig. 5c,e). Other organ age gaps showed no consistent association with network MC (Supplementary Fig. 7). In an exploratory analysis of the wider cohort with accelerometry and baseline proteomics (n ≈ 10,000), older-appearing organs correlated weakly with later sleep: a higher kidney age gap with longer workday sleep (r = 0.042, P = 1.2 × 10⁻⁴), and higher pulmonary, oesophageal, pancreatic, cardiac, male reproductive and proteomic brain age gaps with more irregular sleep (r = 0.031–0.053, P = 1.8 × 10⁻⁴ to 2.1 × 10⁻³). Polygenic scores for depression (β = −0.023 per s.d., 95% CI −0.033 to −0.013, q = 3 × 10⁻⁴), PTSD (β = −0.022, 95% CI −0.032 to −0.012, q = 7 × 10⁻⁴) and insomnia (β = −0.019, 95% CI −0.029 to −0.009, q = 0.010) were associated with lower visual MC (n = 29,575). Scores for PTSD (β = 0.048, 95% CI 0.031 to 0.066, q = 7 × 10⁻⁶), depression (β = 0.041, 95% CI 0.023 to 0.058, q = 3 × 10⁻⁴), problematic alcohol use (β = 0.039, 95% CI 0.021 to 0.057, q = 9 × 10⁻⁴) and alcohol-use disorder (β = 0.032, 95% CI 0.014 to 0.050, q = 0.012) were associated with more irregular sleep (n = 12,212; FDR pooled across secondary tests; Fig. 5d and Supplementary Fig. 8a,b).^30–32^ No polygenic score × sleep interaction survived correction (Supplementary Fig. 8c).

**Fig. 5.**
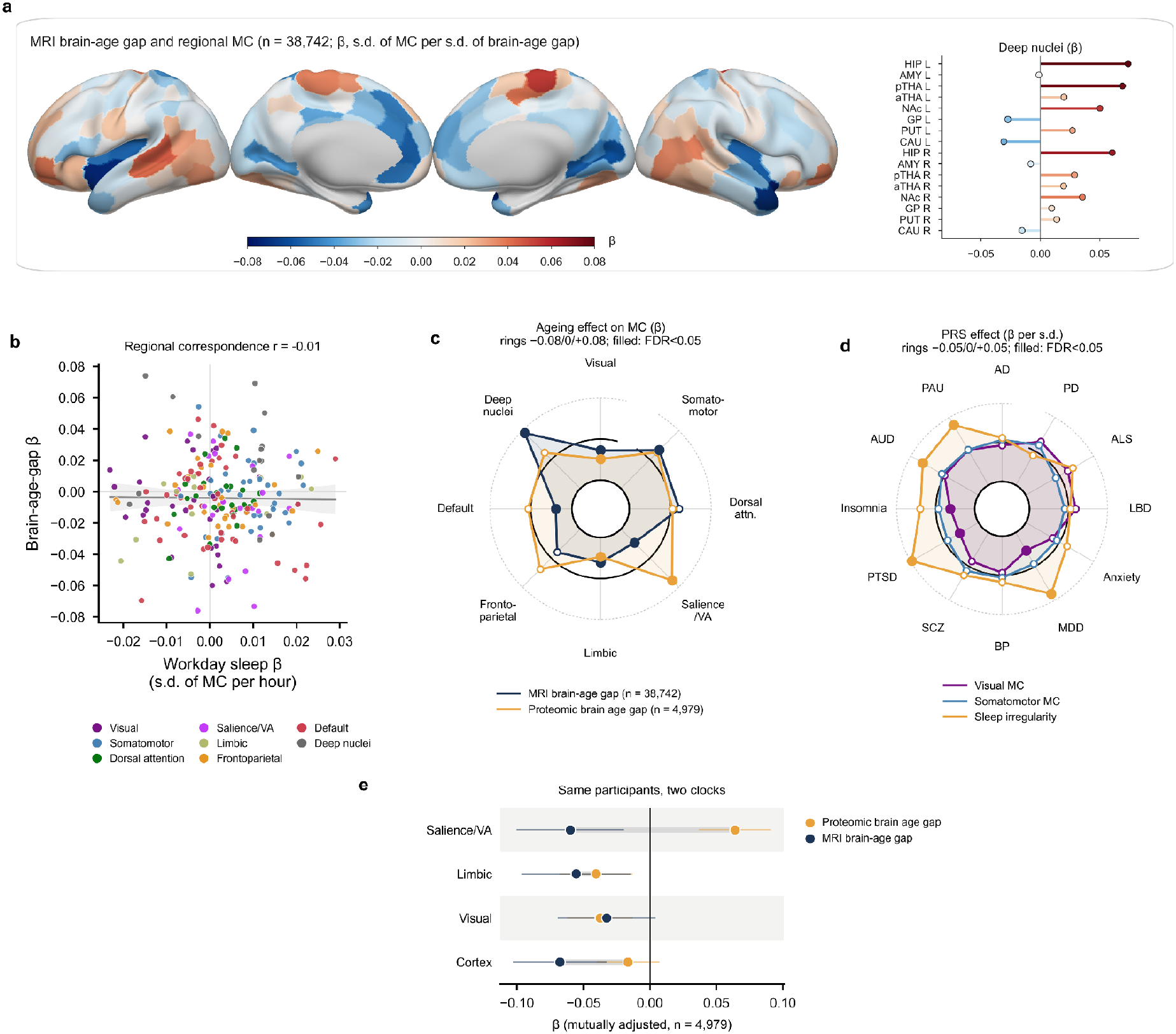
| Biological context. **a**, Regional MC associated with MRI brain-age gap (n = 38,742). **b**, Correspondence of the sleep and brain-age maps (overlapping samples). **c**, Radar of network MC associations (β, s.d. of MC per s.d. of age gap) with MRI brain-age gap (n = 38,742) and proteomic brain-age gap (n = 4,979); bold ring β = 0, dotted rings ±0.08; filled markers, FDR < 0.05. **d**, Radar of polygenic score associations (β per s.d. of PRS) with visual and somatomotor MC (n = 29,575) and sleep irregularity (n = 12,212); rings ±0.05; filled markers, FDR pooled across secondary tests < 0.05. **e**, Mutually adjusted proteomic and MRI brain-age gaps and network MC in the same participants (n = 4,979); grey bars connect the two estimates for each network. Colour bar in **a**: β.

### The signature relates to daily activity and body composition rather than cognition

Finally, we asked which everyday phenotypes travel with the signature, using a hypothesis-free scan with replication in the locked test set. We scanned 1,226 imaging-visit phenotypes, following recent population-neuroscience work (Fig. 6a,b).^33,34^

**Fig. 6.**
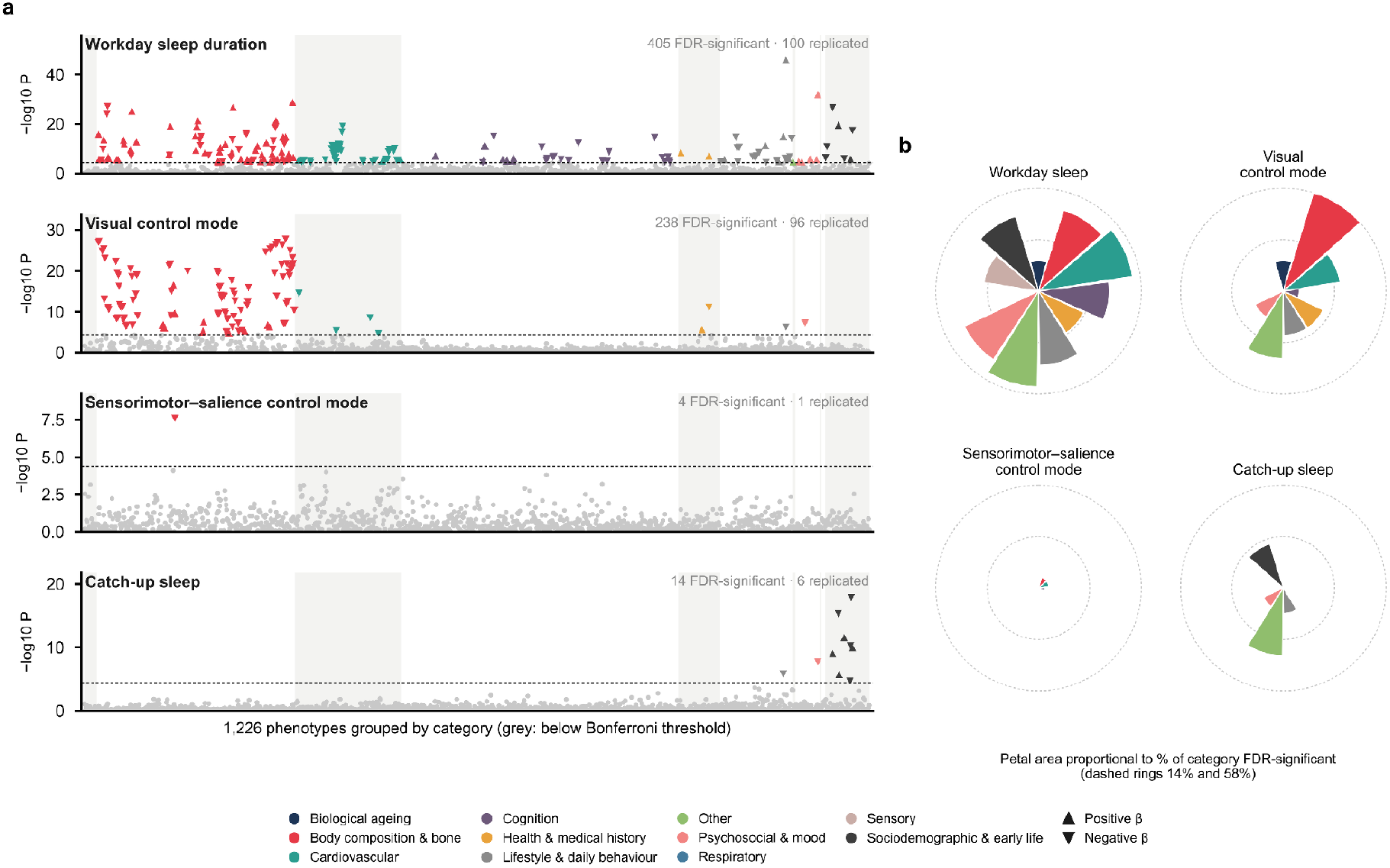
| Functional significance. **a**, Manhattan plots of 1,226 phenotypes against workday sleep, the visual and sensorimotor–salience control modes and catch-up sleep; dashed line, Bonferroni threshold; phenotypes below the threshold in grey; headers give FDR-significant and replicated counts. **b**, Nightingale rose diagrams: for each exposure, each petal is a phenotype category and its area is proportional to the percentage of phenotypes in that category with FDR < 0.05 (dashed rings give reference percentages).

Workday sleep had the broadest profile, with 405 FDR-significant phenotypes; 100 of 137 discovery-Bonferroni hits replicated in the test set. It was associated with more television time (β = 0.115, P = 2 × 10⁻⁴⁶), lower fluid intelligence (β = −0.065, P = 1 × 10⁻¹⁵), a higher trunk fat percentage (β = 0.060, P = 2 × 10⁻¹⁵) and lower fat-free mass (β = −0.018, P = 3 × 10⁻⁴). This suggests that in older adults, longer accelerometer-classified sleep partly marks a less active lifestyle. For the visual control mode, 238 phenotypes were FDR-significant, and 96 of 103 discovery hits replicated. They were dominated by leaner trunk and abdominal composition (for example, trunk tissue fat fraction β = −0.096, P = 2 × 10⁻²⁸; abdominal subcutaneous adipose tissue β = −0.093, P = 6 × 10⁻²⁸) and more physical activity. The sensorimotor–salience mode was largely phenome-silent (4 FDR-significant phenotypes, 1 replicated). For catch-up sleep, associations almost exclusively reflected the working week, including less shift work (β = −0.113, P = 1 × 10⁻¹⁸), employment status (β = −0.064, P = 5 × 10⁻¹⁶), longer working weeks (β = 0.077, P = 1 × 10⁻¹⁰) and commuting (β = 0.088, P = 3 × 10⁻¹²).

Because a phenome scan cannot separate sleep from brain organisation, we then asked whether the sleep–MC association survives adjustment for its strongest phenotypic correlates and which edges of a sleep → MC → phenotype path are supported. Adjusting the primary model for waist circumference, trunk fat, television time and fluid intelligence left the sleep–MC associations essentially unchanged (somatomotor β = 0.025, 95% CI 0.012 to 0.038, P = 1 × 10⁻⁴; visual β = −0.015, 95% CI −0.027 to −0.004, P = 0.010). We then tested each edge of a sleep → structural controllability → phenotype path model separately, rather than requiring a significant indirect effect (Fig. 7). Longer workday sleep was associated with lower visual MC (a path, β = −0.025, 95% CI −0.041 to −0.009, q = 0.008) and higher sensorimotor–salience MC (β = 0.028, 95% CI 0.012 to 0.044, q = 0.004), as was weekend sleep (β = −0.023, q = 0.012; β = 0.019, q = 0.036; n = 14,673); catch-up sleep was not (P = 0.53 and 0.77). Adjusting for sleep, higher visual MC was associated with lower trunk fat fraction (b path, β = −0.083, 95% CI −0.099 to −0.067, n = 15,065) and waist circumference (β = −0.083, 95% CI −0.099 to −0.067, n = 15,299), both with q = 2 × 10⁻²³ and replicated in the discovery and locked test sets separately, and with less television time (β = −0.034, 95% CI −0.049 to −0.018, q = 3 × 10⁻⁴), more physical activity (β = 0.025, 95% CI 0.008 to 0.041, q = 0.032) and faster walking pace (β = 0.021, 95% CI 0.006 to 0.037, q = 0.045), which did not replicate in both sets (Fig. 7a). No sensorimotor–salience path to any phenotype survived correction. Sleep itself was associated with most phenotypes, including lower cognitive scores, more television time and higher trunk fat (Fig. 7b). In the test set, the signature was not associated with fluid intelligence or brain-age gap (Fig. 7c).

**Fig. 7.**
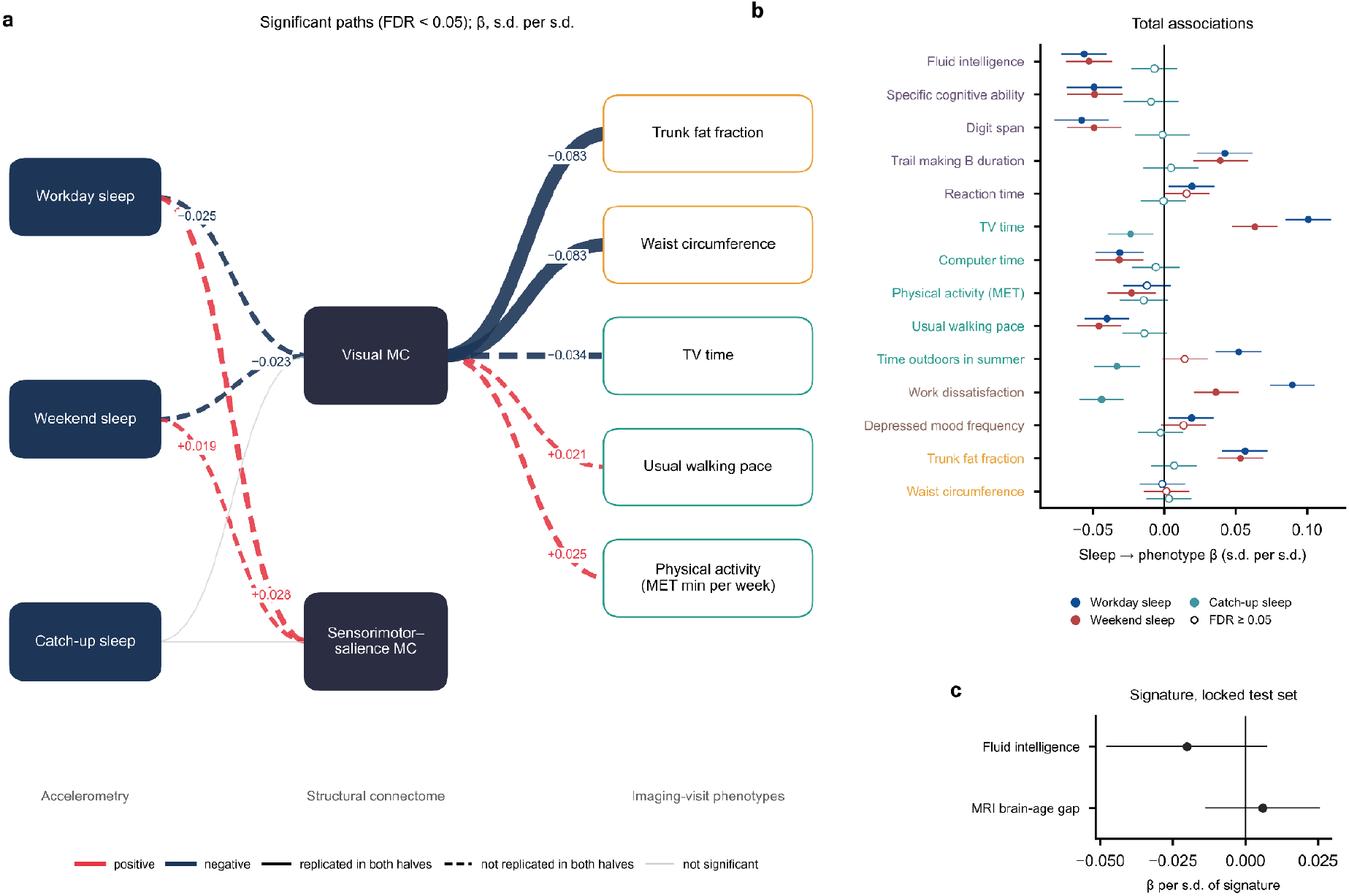
| Sleep, structural controllability and daily behaviour. **a**, Path diagram showing edges that survived FDR correction. Left, accelerometer sleep; middle, network MC modes; right, phenotypes with at least one significant MC edge (box colour, phenotype domain). Sleep → MC edges (a paths) and MC → phenotype edges adjusted for sleep (b paths, from the workday-sleep model) are labelled with standardised β (s.d. per s.d.); line width is proportional to |β|, red positive and blue negative; solid lines replicated in both the discovery and the locked test set, dashed lines did not; grey, not significant. **b**, Total associations of each sleep measure with 14 phenotypes (β with 95% CI; filled, FDR < 0.05; label colour, domain). **c**, Discovery-defined signature and phenotypes in the locked test set (n = 4,382).

## Discussion

Each additional hour of habitual workday sleep was associated with 0.025 s.d. higher somatomotor and 0.018 s.d. lower visual modal controllability, a pattern that reproduced in a locked test set and was absent for weekend catch-up sleep. Because MC in this connectome is an exact function of a region’s share of squared connectivity, the finding is best read as a redistribution of connection weight between sensory systems: more absolute connectivity of visual cortex and a smaller somatomotor share of the whole-brain total, with no significant change in total connectivity. These results address the questions raised in the Introduction: the association is specific to habitual duration rather than to weekend recovery, it reflects a redistribution of connection weight rather than a change in total connectivity, and it is distinct from brain-ageing patterns. We propose, as a hypothesis for future work rather than an established principle, that habitual sleep duration is associated with how the adult connectome apportions connection weight among the systems engaged in waking sensorimotor interaction.

At least three interpretations are compatible with the main association. First, use-dependent and local sleep processes^35,36^ could, over years, shape the connections of the systems most engaged during waking. Second, the direction could be reversed or shared: connectome organisation, or traits that influence it such as health, adiposity and physical activity, could influence how long people sleep. Third, the accelerometer sleep class partly captures quiet rest,^37,38^ so part of the association could reflect sedentary time rather than sleep. The data constrain but do not resolve these alternatives. The association was unchanged after adjustment for diagnoses, health, physical activity, adiposity, television time and cognition, and it was positive in inactive, weekend-warrior and regular-exerciser groups, which argues against confounding by these factors alone. Self-reported sleep at the scan reproduced the visual but not the somatomotor association, so the somatomotor component may depend partly on how sleep is measured. Because sleep and imaging were measured once, the data cannot distinguish the first two interpretations.

Sleep irregularity showed no FDR-significant network association in the full sample, although it was associated with the discovery-defined signature in the test set (β = 0.073 per hour, P = 0.005); its regional maps did not reproduce across split halves. The null result for weekend catch-up sleep also admits more than one reading. Catch-up sleep may simply not relate to connectome organisation, or it may be too noisy a measure to detect an association. Two observations favour the first reading: its regional maps did not reproduce across split halves, and in the phenome scan it tracked the working week (shift work, employment, working hours and commuting) rather than health. Within this cohort, therefore, we found no evidence that weekend recovery is associated with the organisation linked to habitual duration, although we cannot exclude effects in people with severe chronic restriction.

The comparison with ageing separates the signature from an older-appearing brain: the sleep map did not resemble the brain-age map, and the two brain clocks related to salience/ventral-attention MC in opposite directions when entered together. This divergence could reflect different biology captured by plasma proteins and by brain structure, or the nine-year gap between the proteomic and imaging measurements; our data cannot distinguish these possibilities. Genetic liability to depression, PTSD and insomnia was associated with lower visual MC and with more irregular sleep, which is consistent with shared genetic influences on sleep regularity and visual-system wiring, but these are associations at the population level and do not show that genetic risk acts through the connectome. Throughout, we treat spatial correspondence between maps, association across participants, prediction of phenotypes and causation as separate claims; the present data support only the first two.

The phenome scan and the path analysis place the signature in a behavioural context. Higher visual MC was associated with lower trunk fat and waist circumference in both the discovery and the locked test set, whereas the edges to television time, physical activity and walking pace did not replicate. The path analysis tests each edge separately and does not establish mediation: a sleep → MC edge and an MC → phenotype edge can coexist without any causal chain, and the indirect effects were small. The finding is therefore best stated as an association between habitual sleep, sensory-system connection weight and body composition, which future longitudinal and intervention studies could test directly.

### Limitations

First, the data are cross-sectional and cannot show within-person change with sleep or ageing; accelerometry preceded imaging by a median of 3.3 years. Second, precomputed controllability had been derived from unnormalised streamline counts, so average controllability, spectral normalisation and transition energies could not be computed, and MC here is a model-based quantity, not a measure of metabolic energy or of dynamics. Third, we tested independence from regional connection strength and total connectivity but not from inter-regional distance or the cortical hierarchy. Fourth, participant identifiers were recovered by alignment with high but not perfect accuracy, although estimates were unchanged in unambiguously mapped participants. Fifth, effects were small (about 0.02 to 0.05 s.d. per hour, E-values about 1.2), the visual association was not significant without the largest imaging site, and replication was internal to one cohort of largely healthy, older volunteers. These features make the result a well-validated association, not a causal mechanism.

In summary, habitual workday sleep, but not weekend catch-up sleep, is associated with a reproducible redistribution of structural connection weight from somatomotor towards visual cortex that is distinct from brain ageing and travels with body composition.

## Methods

### Participants

We studied UK Biobank participants (application 619657) with a diffusion connectome at the first imaging visit (field 31024) and valid accelerometry (good wear-time and calibration flags).^16,18^ Participants with workday or weekend sleep outside 3–14 h or extreme controllability (|z| > 5) were excluded, leaving n = 15,799. Connectome and polygenic data from a second application were linked through the official crosswalk and the identifiers embedded in the 31024 file names (100% agreement). UK Biobank has ethical approval from the North West Multi-centre Research Ethics Committee, and all participants gave informed consent.

### Recovery of connectome identifiers

Connectomes were derived from the UK Biobank diffusion pipeline.^19,39^ The precomputed controllability matrix (216 × 42,838) lacked identifiers. Its columns follow the sorted names of 31024 bulk folders (both imaging visits), and the download contained 1,039 folders absent from the current release. Participants scanned twice give adjacent columns with near-identical profiles, which provided 2,732 anchors. A banded dynamic-programming alignment matched the 41,799 known folders to the columns, first using the anchors and then adding connectome-predicted sex, age and head size trained on unambiguous segments. Out-of-fold accuracy of connectome-predicted sex was 84.8% in exactly mapped segments and 84.6% in aligned segments versus 51.6% for shuffled identifiers, and predicted age correlated with recorded age at r = 0.63 and 0.64 (Supplementary Fig. 2); because sex and age also informed the second alignment pass, all main estimates were repeated in the exactly mapped subset (Supplementary Methods 4).

### Controllability

The stored values satisfied MC_i_(raw) = 1 − d_i_, so d_i_ = Σ◻ A_i_◻² was recovered exactly. We stabilised the system as A/(1 + ǁAǁ_F)^9,14^, where ǁAǁ_F ≥ λ_max, giving MC_i_ = 1 − d_i_/(1 + ǁAǁ_F)². Network MC was averaged over the nodes of the Schaefer 7-network and Tian S1 groupings.^20–22^ Absolute network squared strength, its share of the whole-brain total and the whole-brain total were analysed separately for decomposition.

### Sleep and activity

Workday and weekend sleep were the sums of the 24 hourly sleep-class fractions (fields 40034 and 40038).^18,38^ Catch-up sleep was weekend minus workday sleep. Irregularity was the s.d. of the seven daily sleep totals. Sleep midpoint was the circular centre of the hourly profile, and social jet lag was the weekend minus workday midpoint.^40^ Physical-activity patterns followed the guideline definition: inactive, <150 min per week of moderate-to-vigorous activity (MVPA); weekend warrior, ≥150 min per week with ≥50% of MVPA in 1–2 days; regular, otherwise.^23^

### Statistics

Outcomes were z-scored. Primary models adjusted for age, age², sex, age × sex, site, intracranial volume, diffusion motion, scanner position, accelerometry-to-scan interval, Townsend index, BMI, education, smoking, alcohol, ethnicity and MVPA. The 70/30 site-stratified split (seed 20261016) was fixed before revision analyses, and the test set was used only for replication. The signature score was the discovery-β-weighted sum of residualised regional MC. Map replication was tested against 1,000 within-test permutations. False discovery rate (FDR) correction^41^ was applied across the six sleep exposures and ten network outcomes, within exposure across 216 nodes for regional maps, and jointly across secondary ageing and genetic tests. Health confounding used hospital ICD-10 diagnoses before the scan (field 41270), self-reported conditions (field 20002), self-rated health (field 2178) and long-standing illness (field 2188).

### Robustness, sensitivity and stability

A specification curve^25^ crossed five samples (all; no shift work; free of the five disease groups; imaging-QC subset; ≥ 6.9 accelerometer wear days), five covariate sets (base; full; full + diagnoses and self-rated health; full + total connectivity; full + scan date, scan and accelerometry season), three outcome treatments (z, rank-based inverse normal, 1%/99% winsorised) and two exposure treatments (raw, winsorised), giving 150 models per association. Joint inference used 500 permutations of the exposure within site, applied identically across specifications, with the median β and the number of same-sign specifications with P < 0.05 as test statistics. We also fitted HC3 standard errors, Huber robust regression and median regression, and refitted after removing the top 1% and 5% of Cook’s distance or outcomes with |z| > 3. Effect modification was tested with F tests for exposure × modifier terms. E-values^26^ were computed from the effect per s.d. of sleep, converted to an approximate risk ratio as exp(0.91 d). Stability analyses used 1,000 nonparametric bootstrap resamples of the full regional model, 200 subsamples without replacement at each of nine sample fractions, leave-one-site-out models and the discovery-defined signature recomputed in the locked test set after setting the weights of one network to zero.

### Ageing and genetics

Organ-specific proteomic age gaps were derived from baseline Olink data.^29^ Following ref. 28, we trained ElasticNet^42^ clocks on organ-enriched proteins with 5-fold cross-fitting and took age-, age²- and sex-adjusted residuals. The MRI brain-age gap came from a cortical-thickness model.^27^ In an exploratory analysis not restricted to the imaging sample, Pearson correlations were computed between each of the 19 proteomic age gaps (baseline, 2006–2010) and the six accelerometer sleep measures (2013–2015) in all participants with valid accelerometry and proteomic data (n = 3,655–9,939 per pair). Polygenic scores for 12 disorders were analysed in unrelated Europeans, adjusting for 10 genetic principal components.^32,43^

### Phenome-wide scan and path analysis

Following PHESANT,^34^ all instance-2 integer and continuous fields were extracted, excluding brain-imaging-derived, procedural and covariate-defining fields. UK Biobank missing codes were removed, and fields with fewer than 1,000 observations or more than 95% in one category were dropped. Continuous phenotypes were rank-inverse-normal transformed, binary phenotypes coded 0/1 and ordinal phenotypes analysed as numeric. Proteomic and MRI ageing measures were added, giving 1,226 phenotypes. Models adjusted for demographic and imaging covariates only. A phenotype was considered replicated if it was Bonferroni-significant in discovery and showed the same sign with P < 0.05 in the test set. For the path analysis, sleep, network MC mode (visual MC; sensorimotor–salience MC, the mean of standardised somatomotor and salience MC) and 14 phenotypes covering cognition, daily behaviour, mood and body composition were residualised on demographic and imaging covariates and standardised. Each edge was estimated by ordinary least squares: sleep → MC (a), MC → phenotype adjusting for sleep (b), sleep → phenotype adjusting for MC (direct) and sleep → phenotype (total). FDR was applied within each edge type, and an edge was considered replicated when it had the same sign and P < 0.05 in both the discovery and the locked test set.

## Supplementary methods and results

Detailed descriptions of every derived measure, model, multiple-comparison family and validation analysis are given in Supplementary Methods 1–17. Numerical results for all figures and analyses are given in Supplementary Tables 1–6 and Supplementary Data 1–6 (provided as separate CSV files).

## Reporting summary

Further information on research design is available in the Nature Portfolio Reporting Summary linked to this article.

## Data availability

UK Biobank data are available to approved researchers via https://www.ukbiobank.ac.uk (application 619657). Derived data supporting the figures are provided as Source Data. [TO COMPLETE: return of derived fields to UK Biobank.]

## Code availability

All analysis code (Python 3.9) is provided in the submission and will be deposited at [TO COMPLETE: repository, DOI].

## Acknowledgements

This research was conducted using the UK Biobank Resource under application 619657. [TO COMPLETE: funding sources and grant numbers.]

## Author contributions

[TO COMPLETE, CRediT format, e.g. X.L. conceived the study, performed the analyses and wrote the manuscript; …]

## Supplementary Information

Supplementary Data 1 (separate CSV file). Regional, network and global associations of all sleep measures with modal controllability (minimally and fully adjusted models).

Supplementary Data 2 (separate CSV file). Discovery and locked test set associations (216 nodes, network and global outcomes).

Supplementary Data 3 (separate CSV file). Phenome-wide scan: all 1,226 phenotypes for each exposure.

Supplementary Data 4 (separate CSV file). Path analysis: all sleep → MC → phenotype edges.

Supplementary Data 5 (separate CSV file). Ageing and polygenic-score associations with MC and sleep, with pooled FDR.

Supplementary Data 6 (separate CSV file). Robustness, sensitivity and stability results (specification curve, subgroups, bootstrap stability, subsampling).

### Supplementary Methods

#### 1. Analytic sample and quality control

Of 100,223 UK Biobank participants with first imaging-visit (instance 2) data, 38,870 had a diffusion connectome and 16,161 of these also had wrist accelerometry. We excluded participants without valid accelerometer wear time or calibration (fields 90015 and 90016), with workday or weekend sleep outside 3–14 h, with extreme MC dispersion (|z| > 5) or with missing intracranial volume or sex, leaving 15,799 participants (Supplementary Fig. 1). Before the revision analyses, these participants were split once into a discovery set (70%, n = 11,059) and a locked test set (30%, n = 4,740), stratified by imaging site with a fixed random seed (20261016). The test set was used only to replicate estimates defined in the discovery set; no model, threshold or region was selected with it.

#### 2. Accelerometer sleep and activity measures

Sleep measures were derived from the 24 hourly sleep-class fractions averaged over workdays (field 40034) and weekend days (field 40038) and from the seven daily sleep totals (field 40042). Workday and weekend sleep durations are the sums of the hourly fractions (hours). Catch-up sleep is weekend minus workday sleep. Sleep irregularity is the standard deviation of the seven daily totals. Sleep midpoint is the circular centre of mass of the 24-h workday profile, and social jet lag is the weekend minus workday midpoint wrapped to ±12 h. Weekly moderate-to-vigorous physical activity (MVPA) is the sum of the seven daily MVPA fractions (field 40045) in minutes. Participants were classified as inactive (< 150 min per week), weekend warriors (≥ 150 min per week with ≥ 50% accumulated on the two most active days) or regular exercisers (≥ 150 min per week otherwise). The accelerometry-to-scan interval is the time from the accelerometer start date (field 90010) to the imaging date.

#### 3. Connectome, modal controllability and decomposition

Structural connectomes were the UK Biobank diffusion-MRI connectomes (field 31024) parcellated into 200 Schaefer cortical regions (7-network assignment) and 16 Tian S1 deep nuclei. The precomputed controllability files contained, for each participant, MC_i_(raw) = 1 − d_i_, where d_i_ = Σ◻ A_i_◻² is the squared connection strength of node i. Because the stabilising normalisation used for the precomputed values could not be recovered, we recomputed MC with the Frobenius normalisation A/(1 + ǁAǁ_F), which guarantees stability because ǁAǁ_F ≥ λ_max, giving MC_i_ = 1 − d_i_/(1 + ǁAǁ_F)². Under this normalisation MC is an exact decreasing function of a node’s share of whole-brain squared connectivity, d_i_/ǁAǁ_F², within each participant (Spearman ρ = −1.000). Network MC is the mean over the nodes of each network; MC dispersion is the standard deviation of MC across all 216 nodes; cortical MC is the mean over the 200 cortical nodes. Whole-brain mean MC is constant by construction and was not analysed. For the decomposition, each network’s MC association was split into its absolute squared connection strength (log mean d_i_), its share of the whole-brain total (mean d_i_/ǁAǁ_F²) and the whole-brain total (log ǁAǁ_F).

#### 4. Recovery and validation of connectome identifiers

The controllability matrix (216 × 42,838) was supplied without participant identifiers. Its columns follow the sorted names of the downloaded diffusion folders (instances 2 and 3), which included 1,039 folders absent from the current data release, so column order alone could not be used. We aligned the 41,799 known folders to the columns by banded dynamic programming. In the first pass, participants scanned at both visits served as anchors: their two scans occupy adjacent columns with highly similar connectome profiles (adjacent-column profile correlation > 0.5; 2,732 column pairs and 2,736 known pairs). Segments between consecutive anchors whose column count equalled the known-folder count were mapped exactly (24,286 participants). In the second pass, ridge and logistic models trained on the exact segments predicted age, sex and head size from the connectome profile and were added to the alignment score to place participants within the remaining segments (17,513 participants). For each participant, the score margin between the chosen column and its neighbours was recorded as a confidence measure. Out-of-fold accuracy of connectome-predicted sex was 84.8% in exactly mapped segments and 84.6% in aligned segments, compared with 51.6% for shuffled identifiers; out-of-fold correlations between predicted and recorded age were 0.63 and 0.64 (Supplementary Fig. 2 and Supplementary Table 1). Because sex and age were also used in the second alignment pass, these figures are an independent validation only for the exactly mapped participants. All main estimates were therefore repeated in the exactly mapped subset (n = 8,799 after sample selection) and in subsets with alignment margins > 2 and > 4 (Fig. 4c). The instance-3 scans were used only as anchors and were genuine repeat scans (median within-person profile correlation r = 0.93 versus r = 0.66 between people; no identical pairs); all analyses used instance-2 data.

#### 5. Covariates and regression models

All outcomes were z-scored across participants. The minimal model adjusted for age, age², sex, age × sex, log intracranial volume, diffusion head motion (field 25922), scanner table and head position (fields 25756–25759), the accelerometry-to-scan interval and imaging site (indicator variables). The full (primary) model additionally adjusted for Townsend deprivation index, body mass index, university degree, previous and current smoking, alcohol intake frequency, White ethnicity and log weekly MVPA. Missing covariate values were median-imputed, with a missing-value indicator added when more than 1% of values were missing. Models were fitted by ordinary least squares; β is the change in MC (s.d.) per unit of exposure (hours for durations, catch-up, irregularity and social jet lag; hours of clock time for the midpoint) and 95% confidence intervals and P values are two-sided.

#### 6. Multiple-comparison families

The false discovery rate (Benjamini–Hochberg) was controlled within pre-defined families: (i) across 216 regional outcomes within each exposure; (ii) across six sleep exposures and ten network and global outcomes for the primary network analysis; (iii) jointly across all secondary ageing and polygenic-score tests (512 tests); (iv) across 72 polygenic score × sleep interaction tests; (v) within exposure across 1,226 phenotypes for the phenome-wide scan; and (vi) within each edge type of the path analysis. Bonferroni thresholds were used to define discovery hits for the phenome-wide replication analysis.

#### 7. Discovery-defined signature and map replication

In the discovery set, the 216 regional β values for workday sleep were used as weights. In the test set, regional MC was residualised on the full covariate set, standardised, and the signature score was computed as the β-weighted sum of residualised regional MC divided by the sum of absolute weights. The signature was then regressed on each sleep measure in the test set. Map replication was the Pearson correlation between discovery and test regional β maps, tested against 1,000 permutations in which workday sleep was shuffled within the test set and the test map re-estimated.

#### 8. Dose–response and catch-up contrasts

Non-linearity was tested by a likelihood-ratio test comparing models with a linear term and with a cubic regression spline (4 degrees of freedom) for workday sleep, both with the full covariate set. Binned estimates (Fig. 2c) are covariate-adjusted means with 95% CIs within workday-sleep bins. Catch-up sleep was analysed continuously and categorically (0–1 h, 1–2 h and > 2 h versus ≤ 0 h), with a linear trend test across categories.

#### 9. Health, disease and imaging-quality analyses

Diagnoses before the imaging date were taken from hospital records (ICD-10, field 41270, with dates from field 41280) and self-report at the imaging visit (field 20002). Groups were sleep disorders (G47, F51; self-report 1123), psychiatric disorders (F10, F2x, F30–F33, F40, F41, F43; self-report 1286, 1287, 1289, 1291), neurological disorders (dementias, Parkinson’s disease, multiple sclerosis, epilepsy, cerebrovascular disease, brain tumours and head injury; F00–F03, G20, G30, G31, G35, G40, G93, I60–I69, C71, D33, S06; corresponding self-report codes) and cardiometabolic disease (hypertension, ischaemic heart disease, atrial fibrillation, heart failure, diabetes and obesity; I10, I11, I20–I25, I48, I50, E10–E14, E66; corresponding self-report codes). Self-rated health (field 2178) and long-standing illness (field 2188) were used as covariates or exclusion criteria. Imaging quality control excluded participants in the top 10% of diffusion head motion or of T1 inverse contrast-to-noise ratio. Each analysis is listed in Supplementary Table 3.

#### 10. Split-half and site analyses

Split-half reproducibility was estimated by randomly dividing the full sample into two halves 100 times, re-estimating the 216 regional β maps in each half and correlating them. Site-specific estimates were obtained by fitting the primary model in each imaging site with at least 100 participants (the Bristol site, n = 20, was not estimable), and leave-one-site-out models excluded each site in turn.

#### 11. Partial least squares

Six sleep measures and 216 regional MC values were residualised on the full covariate set and z-scored. PLS (canonical mode, three latent variables) identified paired sleep and brain patterns. The significance of each latent variable was assessed with 500 permutations of participants in the brain block; loading stability was assessed with 300 bootstrap resamples, expressed as bootstrap ratios (weight divided by its bootstrap standard deviation, with signs aligned to the original solution).

#### 12. Proteomic organ clocks and MRI brain age

Organ age clocks were trained on baseline (instance 0) Olink plasma proteomics using organ-enriched proteins for 11 organ systems (reproductive clocks sex-specific) and 7 gastrointestinal organs; an organismal clock used all proteins. Proteins with more than 20% missing values were removed and the remainder standardised. For each clock, the ElasticNet penalty (l1 ratio 0.5) was chosen by five-fold cross-validation on a random subsample of up to 15,000 participants and then fixed; five-fold cross-fitting gave an out-of-fold predicted age for every participant. Organ age gaps are the residuals of predicted age on age, age² and sex, z-scored (performance in Supplementary Table 4). The MRI brain-age gap is predicted minus chronological age from a cortical-thickness model at the imaging visit.

#### 13. Polygenic scores and gene–environment interactions

Polygenic scores for 12 neurological and psychiatric disorders were analysed in unrelated participants of European ancestry, adjusting for the full covariate set and 10 genetic principal components. Gene–environment interactions were tested by adding a polygenic score × sleep product term to models that included both main effects.

#### 14. Phenome-wide scan

All instance-2 integer and continuous fields were extracted, excluding brain-imaging-derived, procedural and covariate-defining fields. Negative UK Biobank special codes were set to missing. Fields with fewer than 1,000 observations or more than 95% of values in one category were removed. Binary fields were coded 0/1, fields with 3–10 levels were analysed as ordinal numeric variables and the remaining continuous fields were rank-based inverse-normal transformed. Each phenotype was regressed on the exposure with the minimal (demographic and imaging) covariate set, so that lifestyle phenotypes were not adjusted away. A discovery hit was Bonferroni-significant in the discovery set; it was considered replicated if it had the same sign and P < 0.05 in the test set.

#### 15. Path analysis

Sleep, MC modes (visual MC; sensorimotor–salience MC, the standardised mean of somatomotor and salience MC) and 14 phenotypes were residualised on the minimal covariate set plus Townsend index, university degree and ethnicity, and standardised. For each sleep measure, mode and phenotype, ordinary least-squares models estimated the sleep → MC edge (a), the MC → phenotype edge adjusted for sleep (b), the sleep → phenotype edge adjusted for MC (direct) and the total sleep → phenotype association. An edge was considered replicated if it had the same sign and P < 0.05 in both the discovery and the test set. No indirect (mediated) effect is claimed.

#### 16. Robustness, sensitivity and stability analyses

The specification curve, permutation inference, robust estimators, influence analyses, additional confounders, effect modification, E-values, bootstrap stability, subsampling, leave-one-site-out and leave-one-network-out analyses are described in Methods. E-values were computed from the effect per s.d. of sleep (d), converted to an approximate risk ratio RR = exp(0.91 × d) and E = RR + √(RR × (RR − 1)), for the point estimate and for the confidence limit closest to the null.

#### 17. Exploratory organ-age and sleep correlations

To use all participants with both measures, the correlations between the 19 proteomic organ age gaps and the six accelerometer sleep measures were computed in the full cohort, not restricted to the imaging sample, using participants with valid accelerometry (wear and calibration flags; workday and weekend sleep 3–14 h) and proteomic data (n = 3,655–9,939 per pair; Pearson correlations, two-sided P values).

---

**Supplementary Fig. 1.**
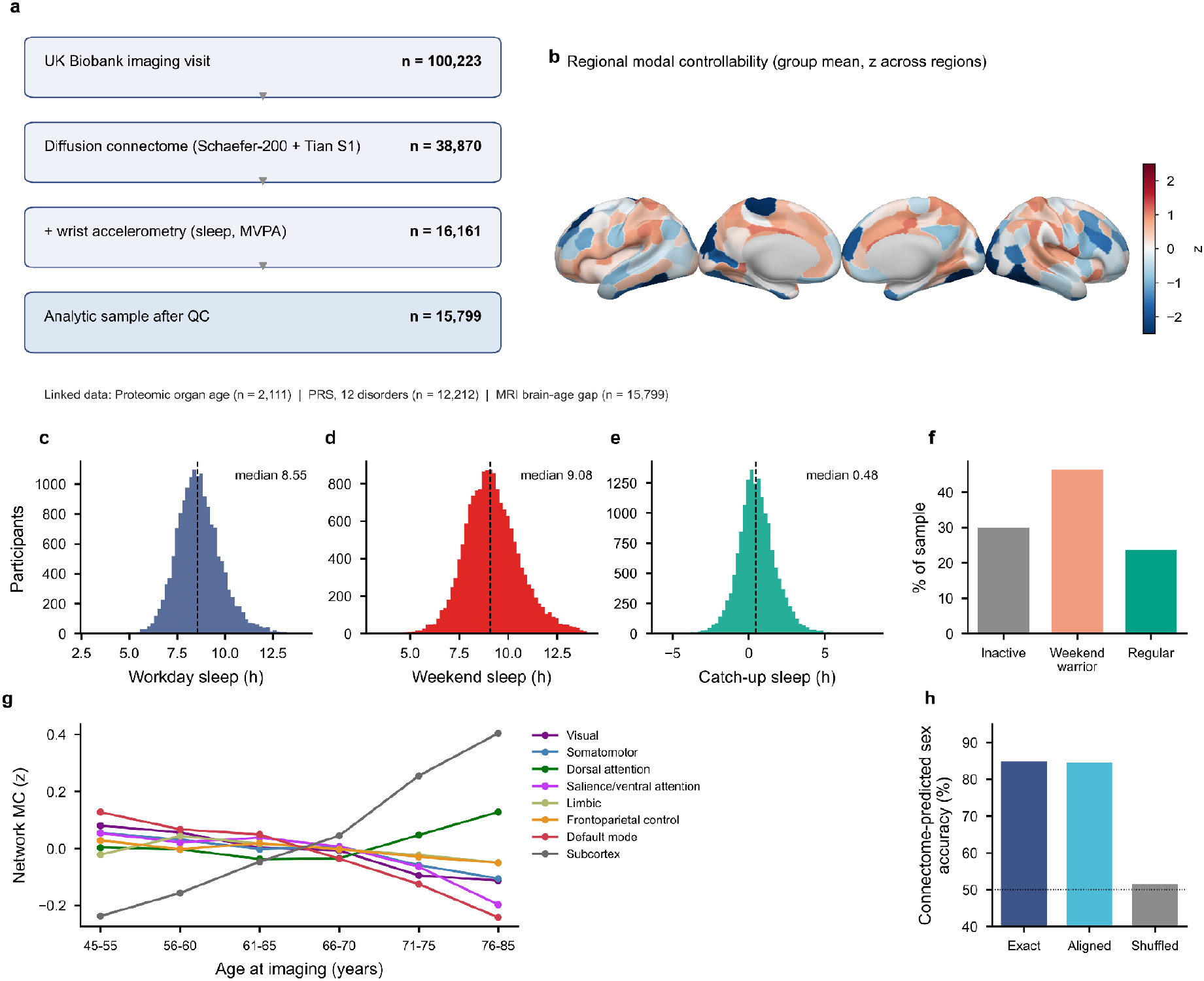
| Study population and data overview. **a**, Participant flow from the UK Biobank imaging visit to the analytic sample. **b**, Group-mean regional modal controllability (MC; z across regions). **c–e**, Distributions of workday, weekend and catch-up sleep (dashed line, median). **f**, Physical-activity patterns. **g**, Network MC by age band (mean z). **h**, Accuracy of connectome-predicted sex for exactly matched, aligned and shuffled identifiers (dotted line, chance).

**Supplementary Fig. 2.**
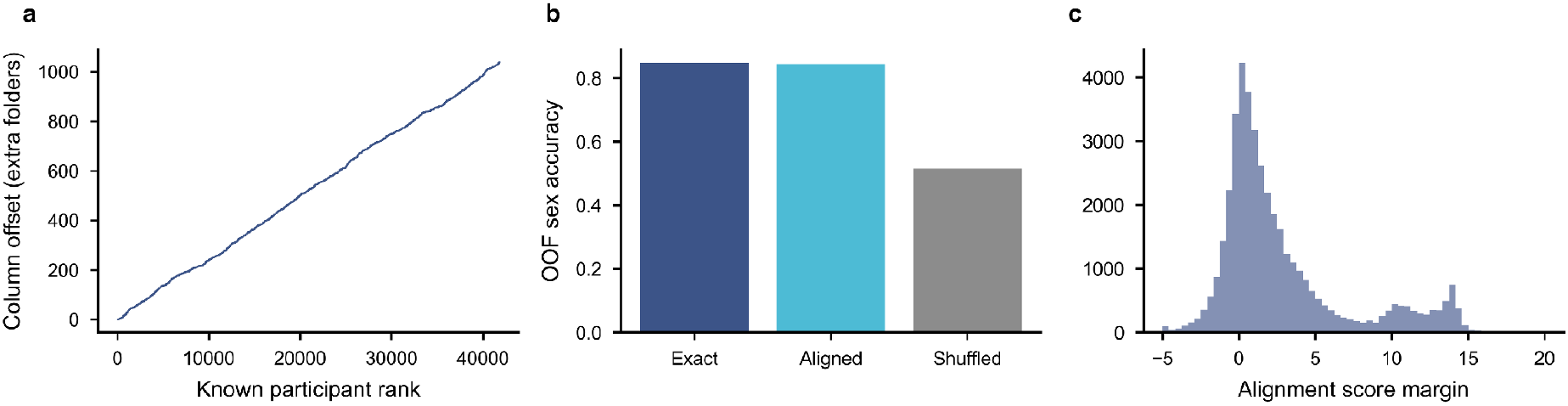
| Recovery of connectome identifiers. **a**, Column offset of each known participant (cumulative number of unmatched folders) along the alignment. **b**, Out-of-fold accuracy of connectome-predicted sex for exactly matched, aligned and shuffled identifiers. **c**, Distribution of alignment score margins.

**Supplementary Fig. 3.**
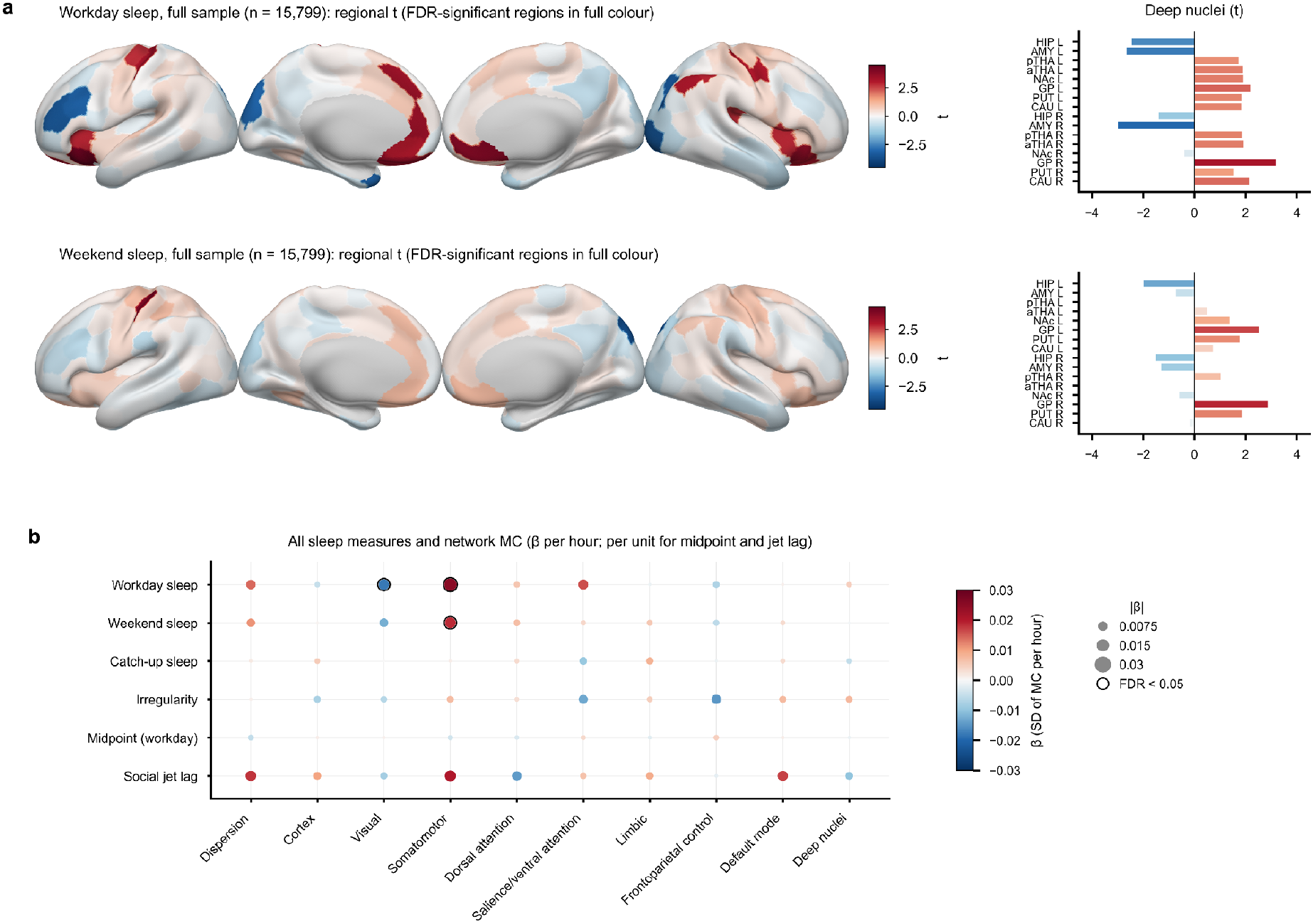
| Full-sample maps and all sleep measures. **a**, Regional t-statistics for workday (top) and weekend (bottom) sleep in the full sample (n = 15,799); FDR-significant regions are shown in full colour and others at 35% intensity; bars show the 16 deep nuclei. **b**, Associations of six sleep measures with network MC. Colour gives β (s.d. of MC per hour; per hour of clock time for midpoint and social jet lag), dot area is proportional to |β| and a black ring marks FDR < 0.05 pooled across exposures and outcomes.

**Supplementary Fig. 4.**
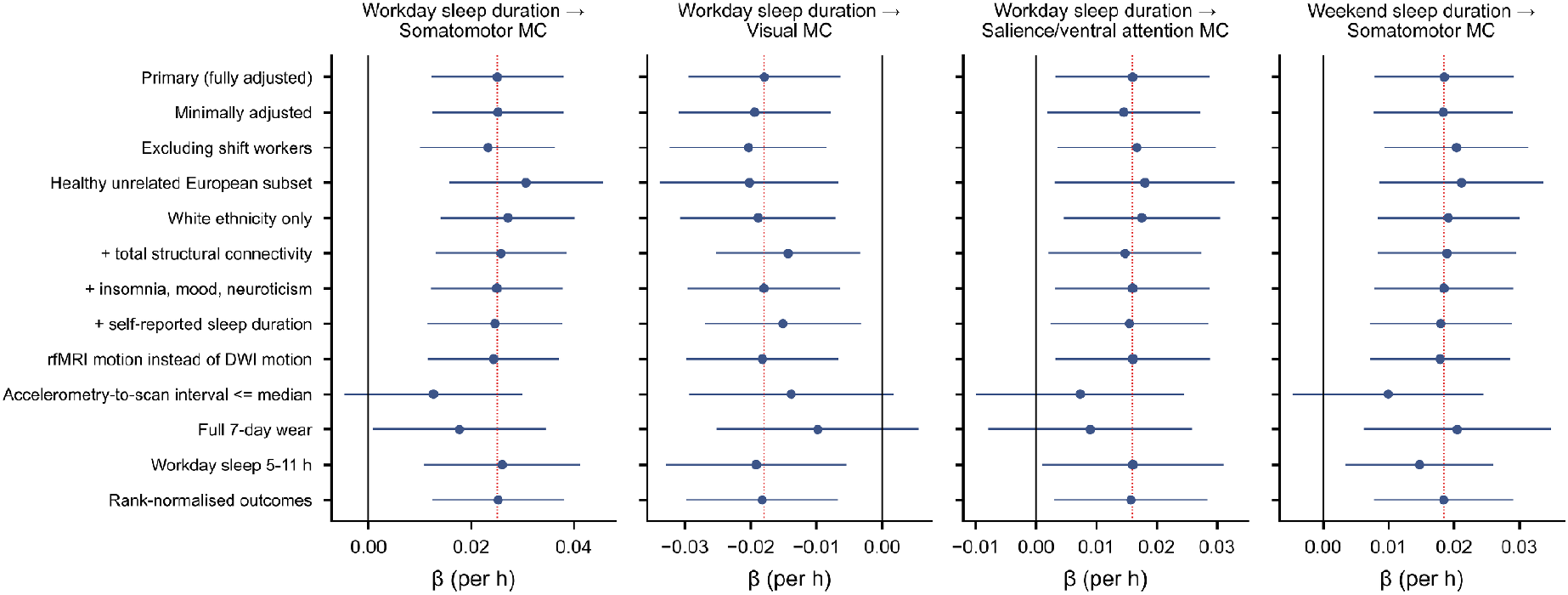
| Covariate and sample sensitivity analyses. Workday-sleep associations with somatomotor, visual and salience/ventral-attention MC, and the weekend-sleep association with somatomotor MC, across 13 model and sample variants (β per hour with 95% CI; red dotted line, primary estimate).

**Supplementary Fig. 5.**
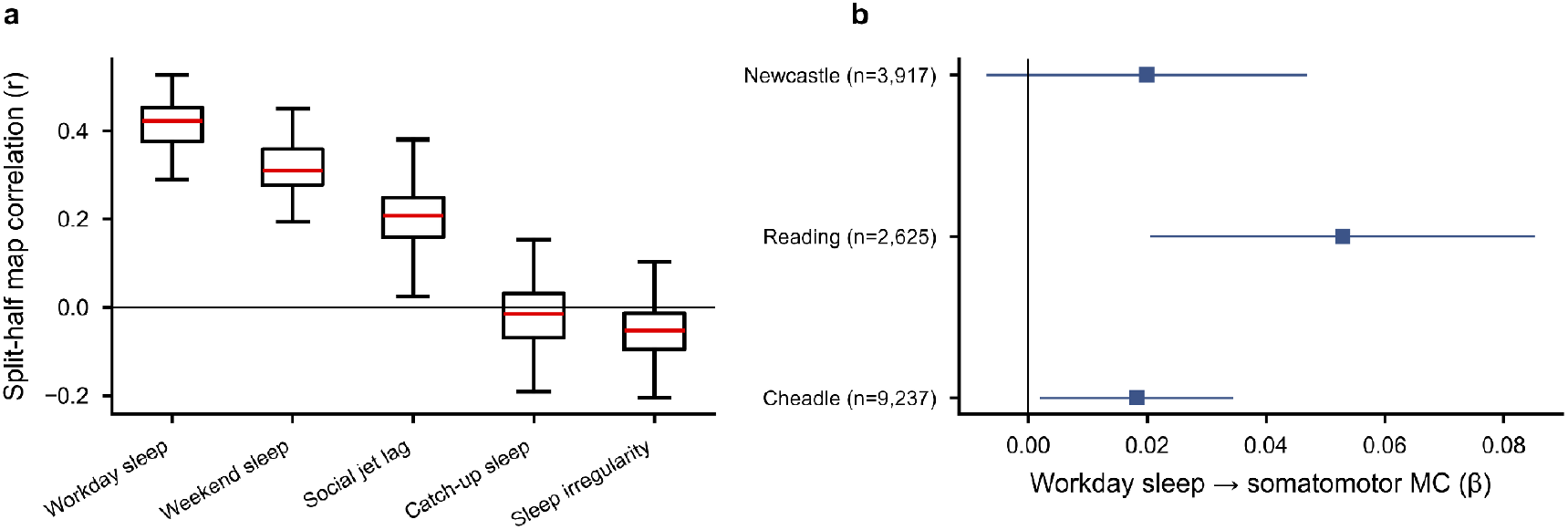
| Split-half and site replication. **a**, Correlation between regional maps estimated in random halves of the sample (100 splits; boxes show the median and interquartile range). **b**, Workday sleep and somatomotor MC within each imaging site with an estimable sample (β per hour with 95% CI).

**Supplementary Fig. 6.**
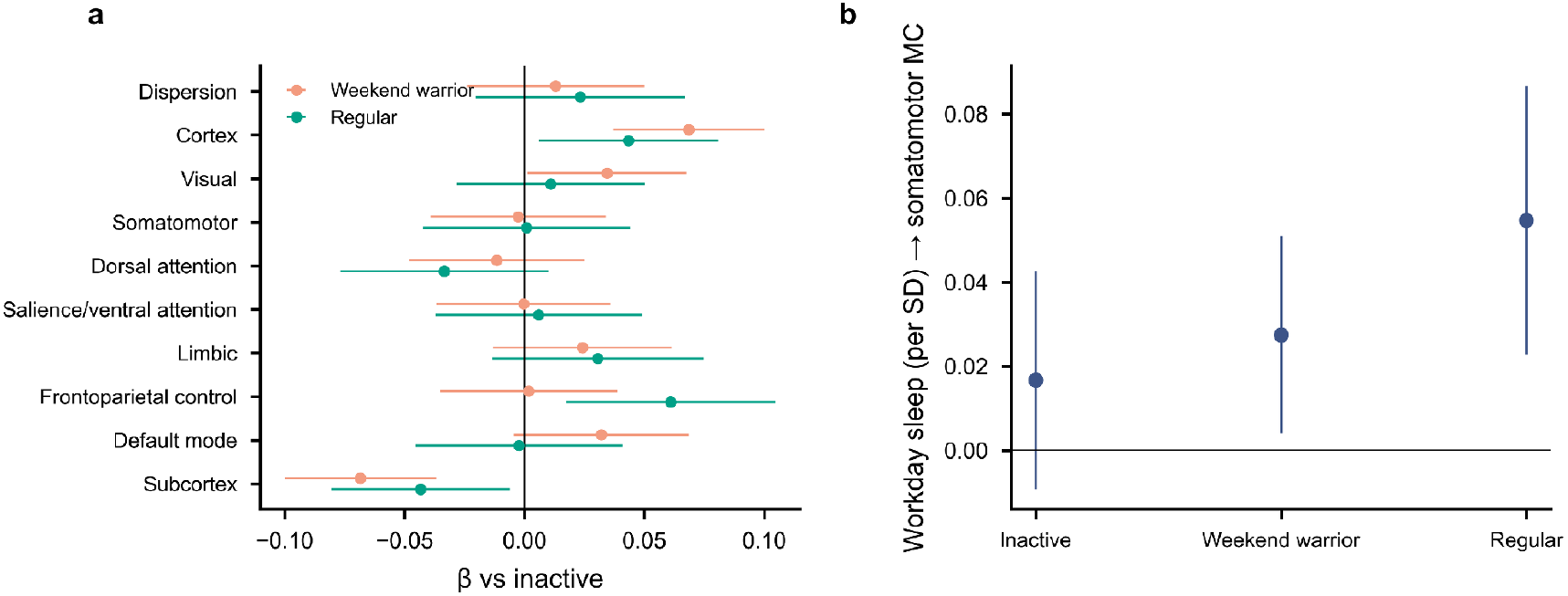
| Physical-activity patterns. **a**, Network MC in weekend warriors and regular exercisers relative to inactive participants (β with 95% CI). **b**, Workday sleep and somatomotor MC within each activity pattern (β per s.d. of sleep with 95% CI).

**Supplementary Fig. 7.**
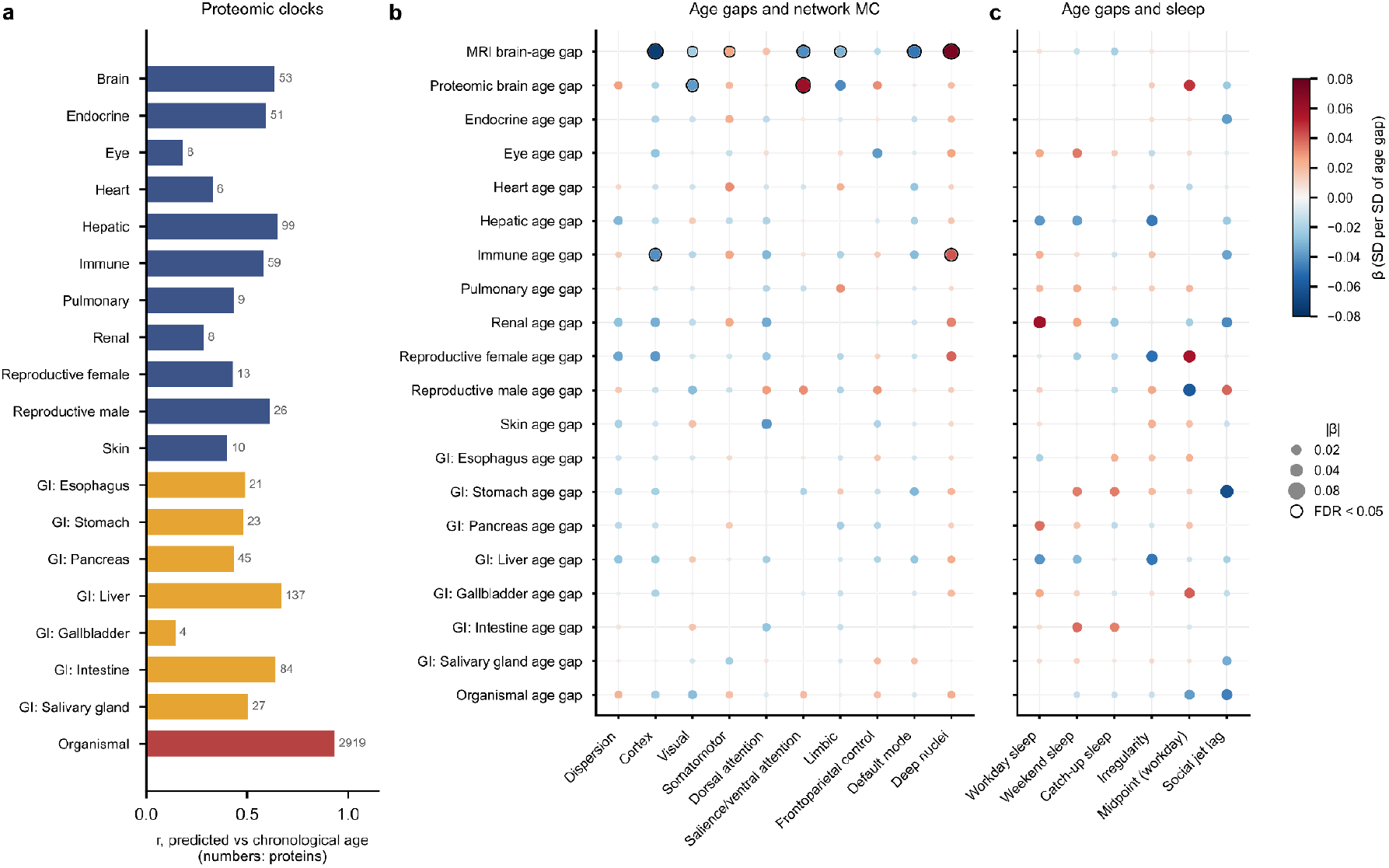
| Proteomic organ ageing. **a**, Performance of the organ-specific proteomic clocks (cross-fitted correlation between predicted and chronological age; numbers give the proteins in each clock). **b**, Associations of MRI and proteomic age gaps with network MC. **c**, Associations of age gaps with sleep measures. In **b** and **c**, colour gives β (s.d. per s.d. of age gap), dot area is proportional to |β| and a black ring marks FDR < 0.05 pooled across all secondary tests.

**Supplementary Fig. 8.**
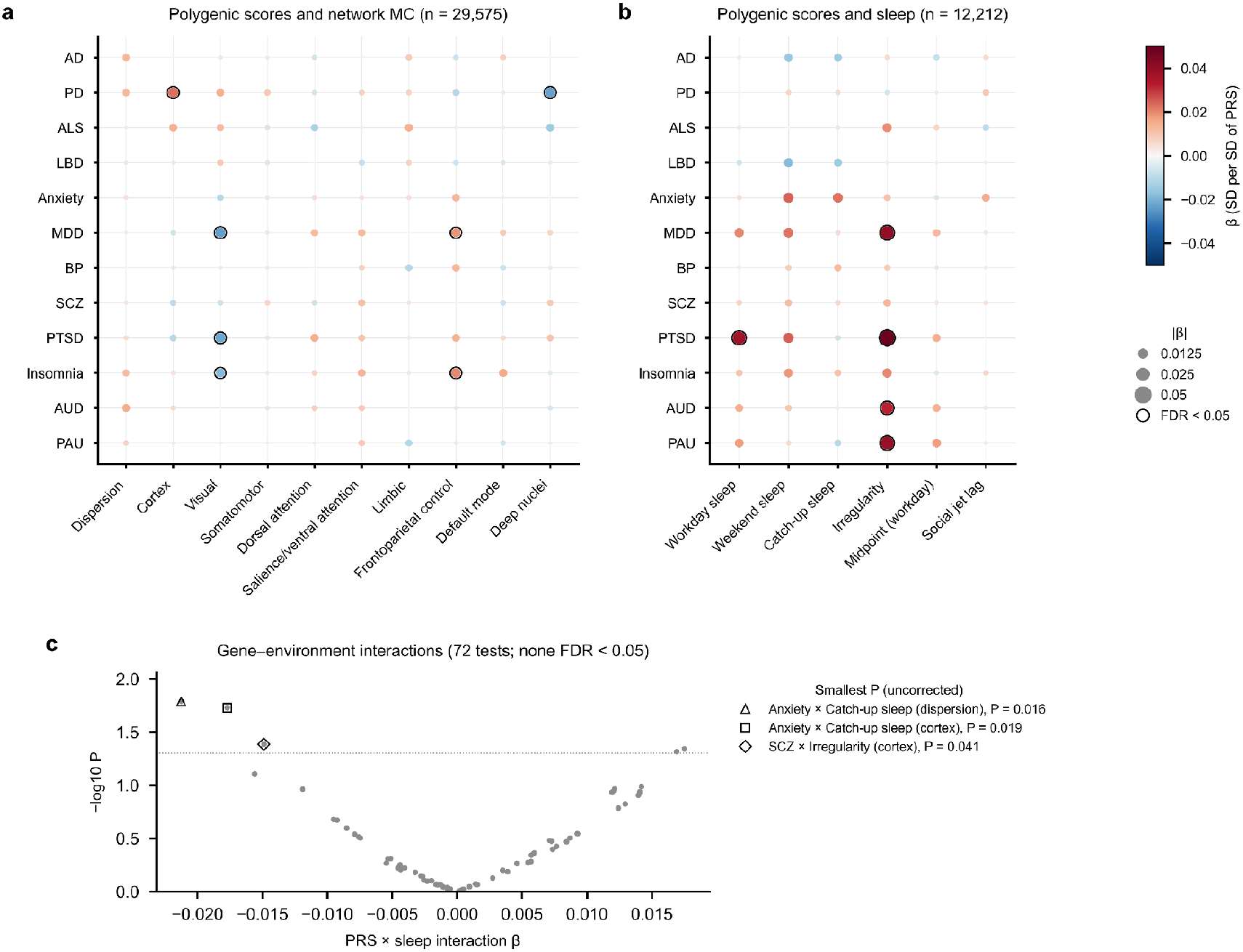
| Polygenic scores and gene–environment interactions. **a**, Polygenic scores for 12 disorders and network MC (n = 29,575). **b**, Polygenic scores and sleep measures (n = 12,212). Colour gives β per s.d. of polygenic score, dot area is proportional to |β| and a black ring marks FDR < 0.05 pooled across secondary tests. **c**, Polygenic score × sleep interactions on MC (72 tests); dotted line, P = 0.05; open symbols mark the three smallest P values, none of which survived FDR correction. AD, Alzheimer’s disease; PD, Parkinson’s disease; ALS, amyotrophic lateral sclerosis; LBD, Lewy body dementia; MDD, major depressive disorder; BP, bipolar disorder; SCZ, schizophrenia; PTSD, post-traumatic stress disorder; AUD, alcohol-use disorder; PAU, problematic alcohol use.

**Supplementary Fig. 9.**
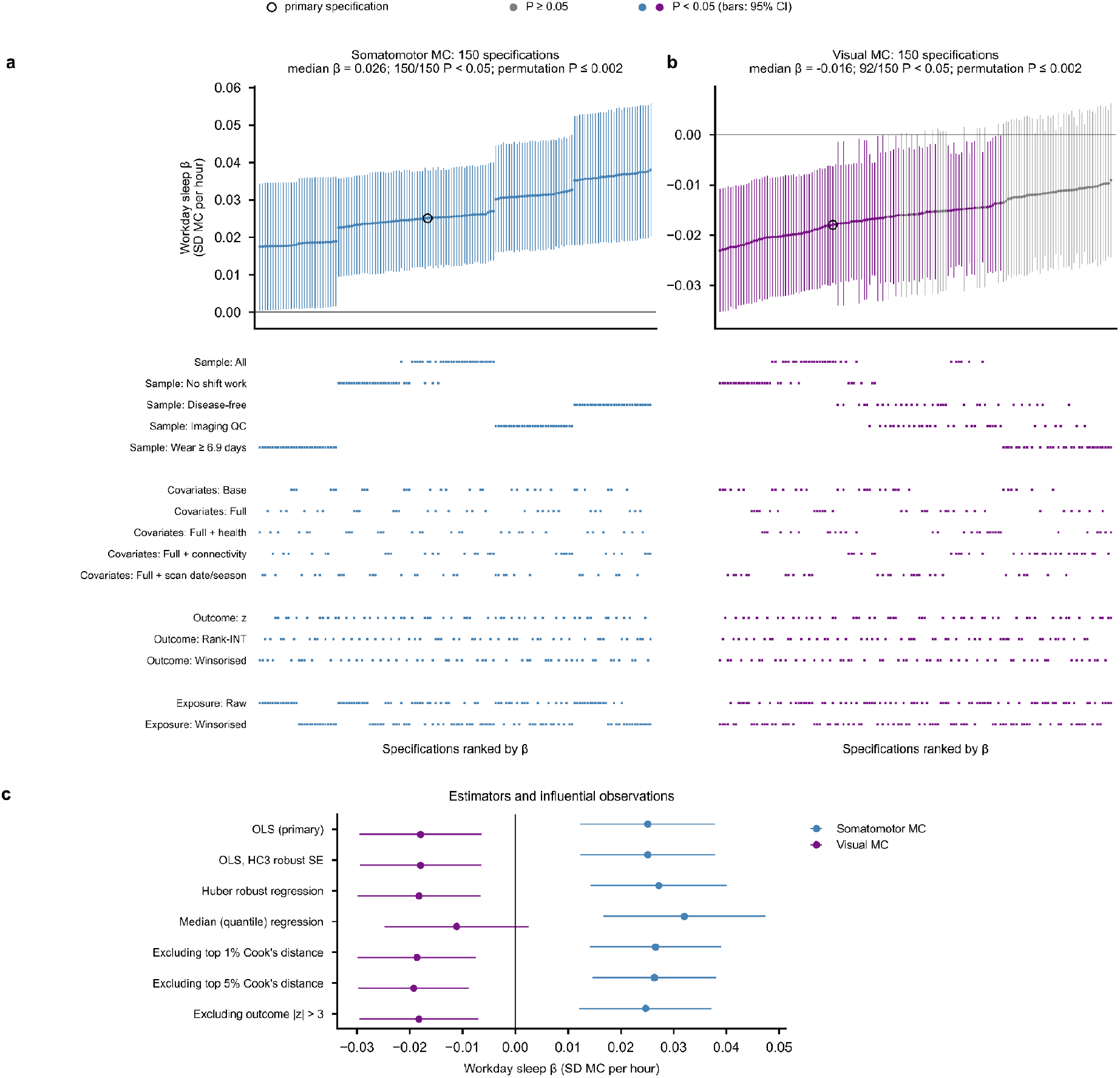
| Robustness: specification curve and estimators. **a,b**, Workday-sleep β (s.d. of MC per hour; 95% CI) for somatomotor (**a**) and visual (**b**) MC across 150 specifications ranked by β; coloured, P < 0.05; open circle, primary specification. The lower panels mark the analytic choice used in each specification. Headers give the median β, the number of specifications with P < 0.05 and the joint permutation P (500 within-site permutations). **c**, Alternative estimators and removal of influential observations.

**Supplementary Fig. 10.**
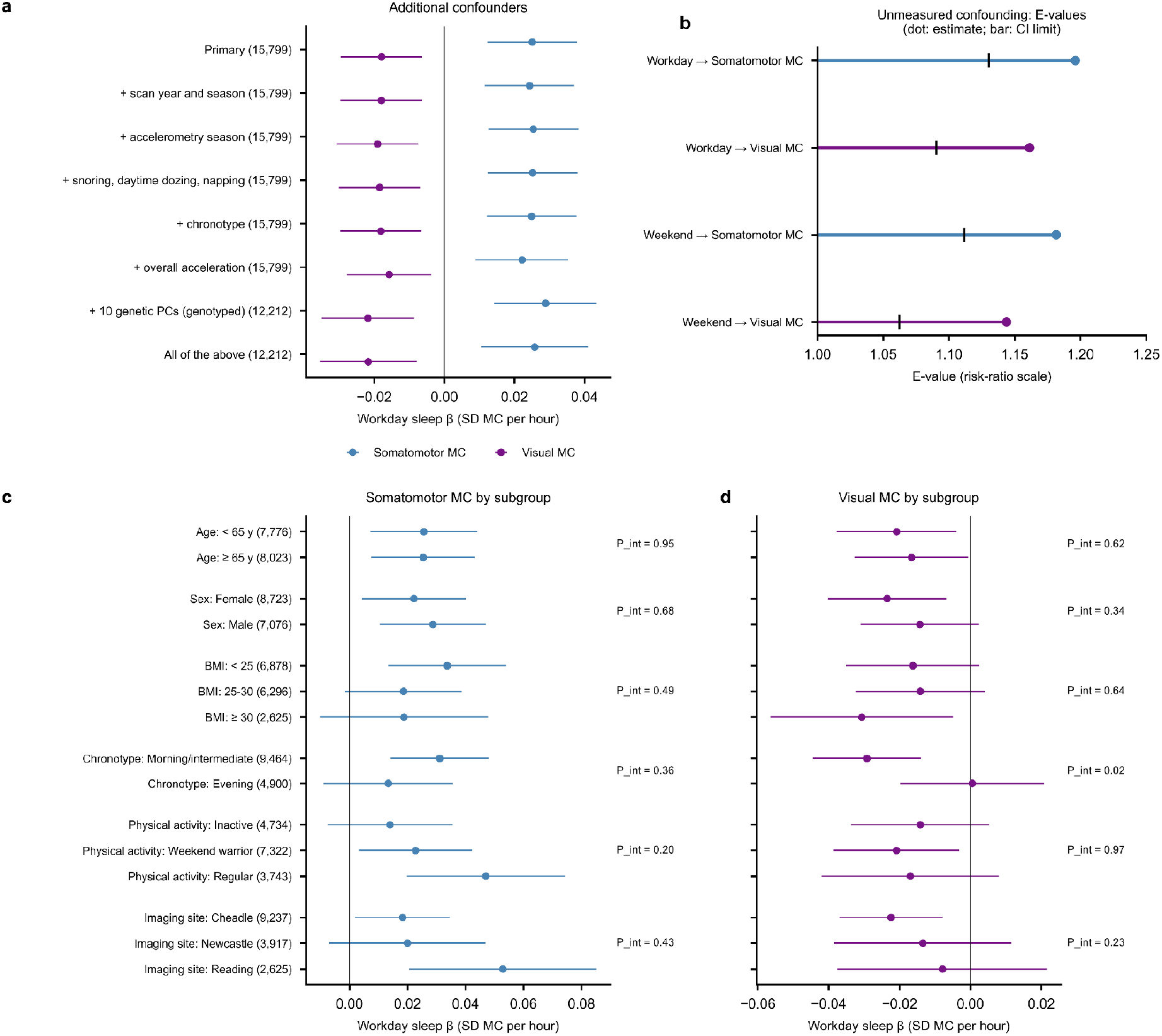
| Sensitivity: confounding and effect modification. **a**, Workday-sleep β after additional adjustment (n in brackets). **b**, E-values for the point estimate (dot) and the confidence limit closest to the null (bar). **c,d**, Workday-sleep β within subgroups for somatomotor (**c**) and visual (**d**) MC; P_int, F test of the exposure × modifier interaction (uncorrected).

**Supplementary Fig. 11.**
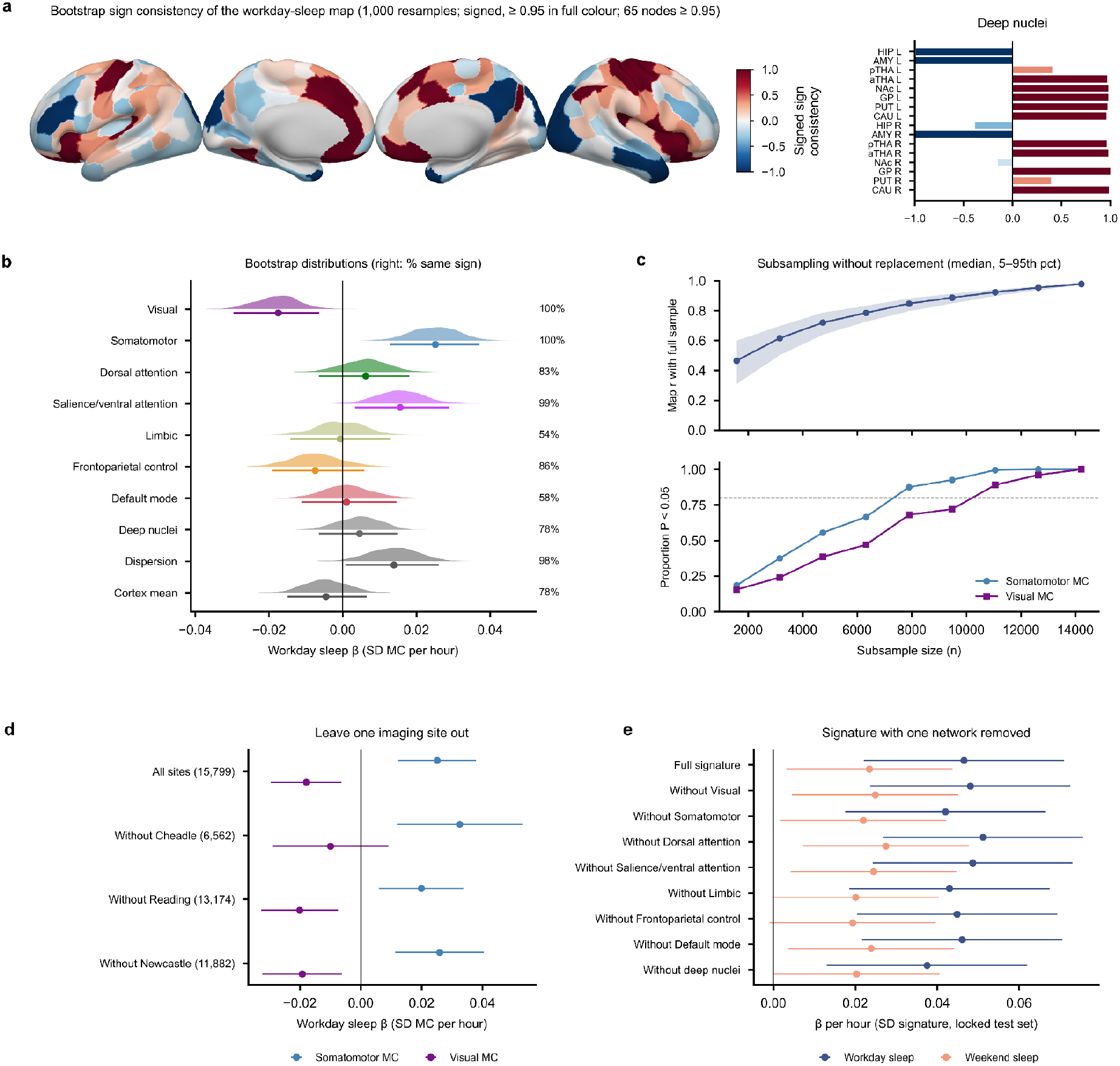
| Stability. **a**, Signed bootstrap sign consistency of the regional workday-sleep map (1,000 resamples): the sign of β multiplied by the proportion of resamples with that sign; nodes ≥ 0.95 are shown at full intensity. **b**, Bootstrap distributions of network β (half-violins; median and 95% interval below); right, percentage of resamples with the full-sample sign. **c**, Subsampling without replacement (200 draws per size): correlation of the subsample map with the full-sample map (median and 5th–95th percentile; top) and proportion of draws with P < 0.05 (bottom; dashed line, 0.8). **d**, Leave-one-site-out estimates. **e**, Discovery-defined signature in the locked test set with the weights of one network set to zero.

**Supplementary Table 1.** | Identifier recovery summary.

| **Statistic** | **Value** |
| --- | --- |
| known folders | 41,799 |
| columns | 42,838 |
| extra columns | 1,039 |
| anchors L | 2,736 |
| anchors C | 2,732 |
| exact | 24,286 |
| pass agree | 32,477 |
| oof sex acc exact | 0.848 |
| age r exact | 0.631 |
| icv r exact | 0.716 |

Alignment of the $216 \times 42,838$ controllability matrix to 41,799 known diffusion folders (Methods).

**Supplementary Table 2.**
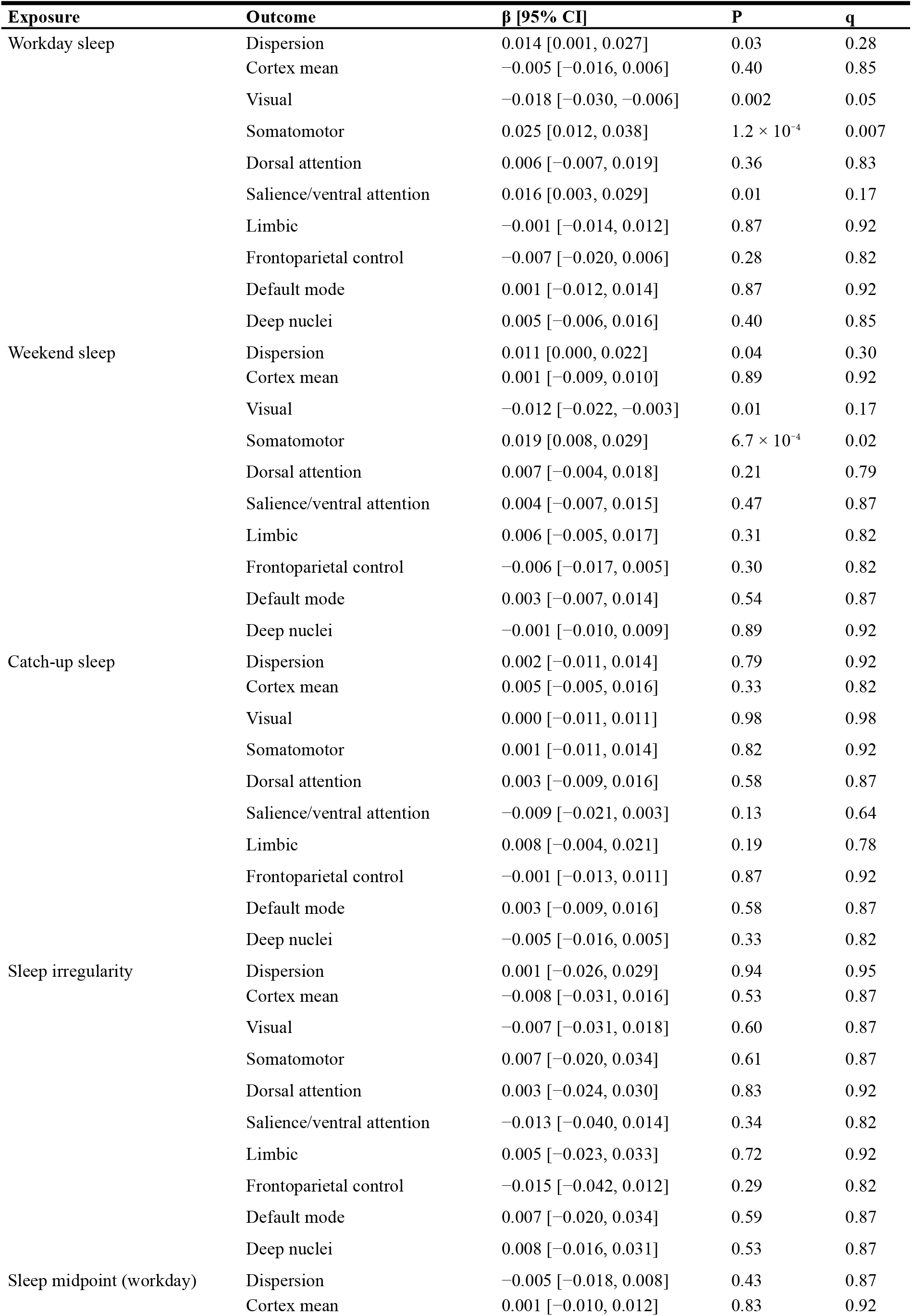
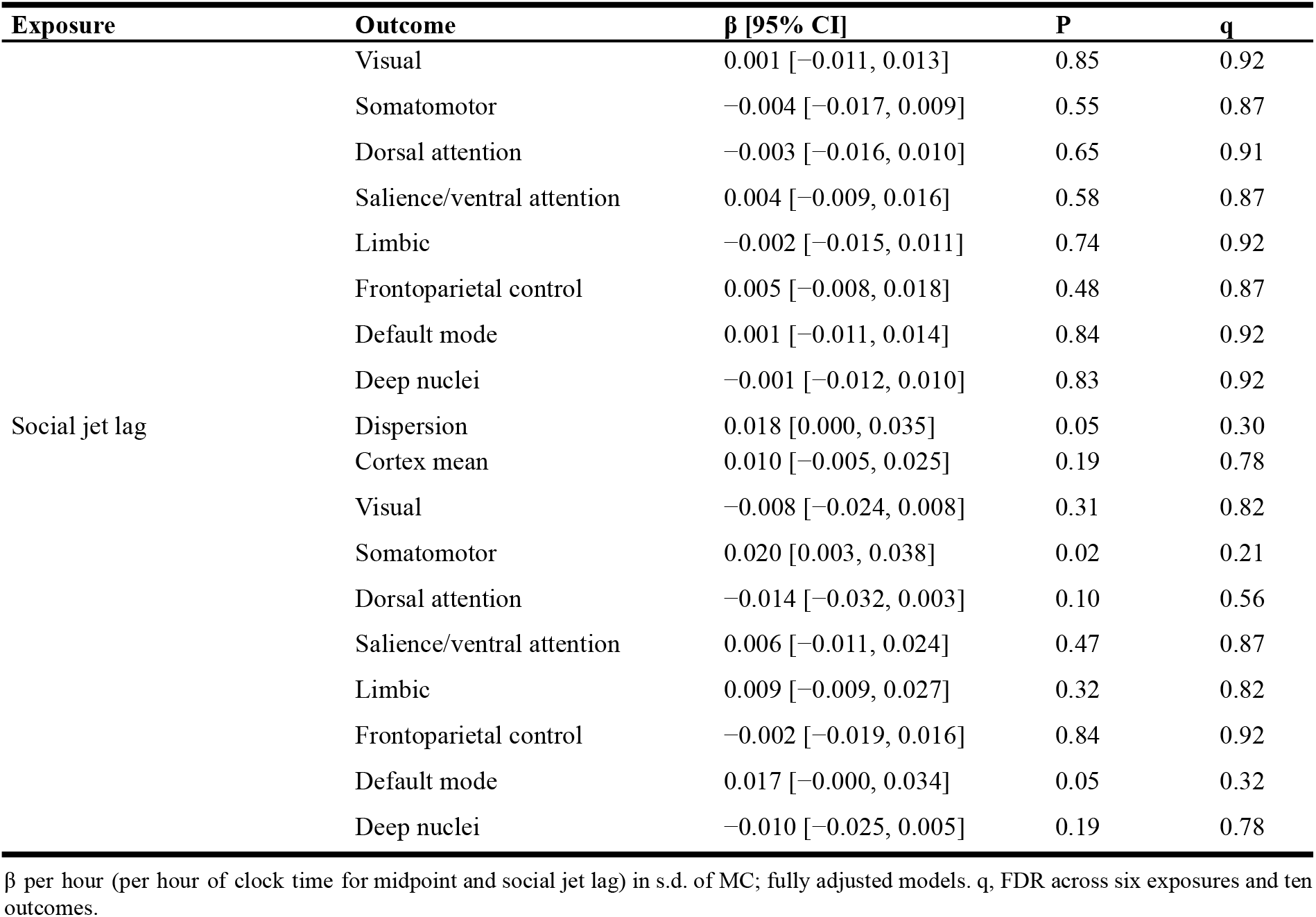
| Sleep measures and network modal controllability (full sample, n = 15,799).

**Supplementary Table 3.** | Health, disease and imaging-quality analyses for workday sleep.

| Analysis | n | Somatomotor MC $\beta$ [95% CI] | Visual MC $\beta$ [95% CI] |
| --- | --- | --- | --- |
| Primary | 15,799 | 0.025 [0.012, 0.038] | -0.018 [-0.030, -0.006] |
| + adjust diagnoses, overall health, long-standing illness | 15,799 | 0.025 [0.012, 0.037] | -0.017 [-0.029, -0.006] |
| Exclude sleep disorders | 15,617 | 0.025 [0.012, 0.038] | -0.017 [-0.029, -0.006] |
| Exclude psychiatric disorders | 14,264 | 0.030 [0.016, 0.043] | -0.020 [-0.032, -0.007] |
| Exclude neurological disorders | 15,284 | 0.025 [0.012, 0.038] | -0.019 [-0.031, -0.007] |
| Exclude cardiometabolic disease | 11,230 | 0.028 [0.012, 0.043] | -0.012 [-0.026, 0.002] |
| Exclude fair/poor self-rated health | 13,215 | 0.029 [0.014, 0.043] | -0.019 [-0.032, -0.006] |
| Disease-free & good health | 8,015 | 0.040 [0.021, 0.058] | -0.018 [-0.035, -0.000] |
| Imaging QC: exclude top-10% motion / T1 CNR | 12,752 | 0.031 [0.017, 0.045] | -0.016 [-0.029, -0.003] |

Diagnoses from hospital records (field 41270) and self-report (field 20002) before the scan; self-rated health (field 2178); long-standing illness (field 2188).

**Supplementary Table 4.** | Proteomic organ clocks.

| **Organ** | **n** | **Proteins** | **r** | **MAE, years** |
| --- | --- | --- | --- | --- |
| Brain | 51,314 | 53 | 0.636 | 5.15 |
| Endocrine | 51,967 | 51 | 0.593 | 5.41 |
| Eye | 44,377 | 8 | 0.181 | 6.85 |
| Heart | 44,729 | 6 | 0.329 | 6.51 |
| Hepatic | 44,873 | 99 | 0.650 | 5.05 |
| Immune | 52,096 | 59 | 0.579 | 5.47 |
| Pulmonary | 52,022 | 9 | 0.432 | 6.16 |
| Renal | 44,577 | 8 | 0.285 | 6.64 |
| Reproductive female | 27,994 | 13 | 0.427 | 6.04 |
| Reproductive male | 20,612 | 26 | 0.611 | 5.40 |
| Skin | 51,826 | 10 | 0.402 | 6.28 |
| GI: Esophagus | 51,614 | 21 | 0.488 | 5.94 |
| GI: Gallbladder | 51,592 | 4 | 0.145 | 6.93 |
| GI: Intestine | 51,495 | 84 | 0.641 | 5.14 |
| GI: Liver | 44,976 | 137 | 0.668 | 4.94 |
| GI: Pancreas | 52,002 | 45 | 0.433 | 6.17 |
| GI: Salivary gland | 51,365 | 27 | 0.503 | 5.88 |
| GI: Stomach | 44,913 | 23 | 0.479 | 5.98 |
| Organismal | 45,968 | 2919 | 0.932 | 2.36 |

ElasticNet clocks trained on organ-enriched Olink proteins with 5-fold cross-fitting; r and MAE are out-of-fold. GI, gastrointestinal.

**Supplementary Table 5.** | Specification-curve summary.

| Exposure | Outcome | Specifications | Median $\beta$ | Range | Same sign, % | P < 0.05, n | Permutation P |
| --- | --- | --- | --- | --- | --- | --- | --- |
| Workday sleep | Somatomotor | 150 | 0.026 | 0.017 to 0.038 | 100 | 150 | 0.002 |
| Workday sleep | Visual | 150 | -0.016 | -0.023 to -0.009 | 100 | 92 | 0.002 |
| Weekend sleep | Somatomotor | 150 | 0.020 | 0.018 to 0.027 | 100 | 150 | 0.002 |
| Weekend sleep | Visual | 150 | -0.013 | -0.018 to -0.007 | 100 | 84 | 0.004 |

Specifications cross five samples, five covariate sets, three outcome and two exposure treatments. Permutation P from 500 within-site permutations applied to all specifications (test statistic: number of same-sign specifications with $P < 0.05$ ); 0.002 is the smallest attainable value.

**Supplementary Table 6.**
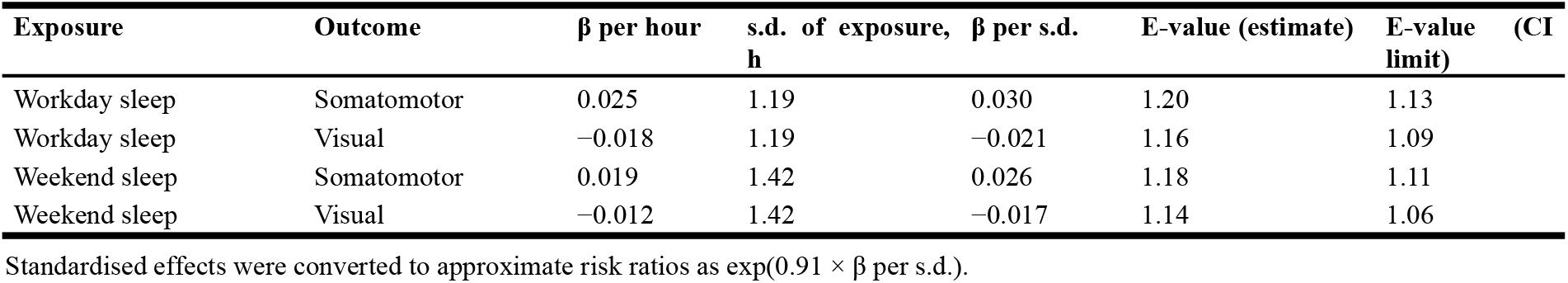
| E-values for unmeasured confounding.

